# Family matters: Linking population growth, kin interactions, and African elephant social groups

**DOI:** 10.1101/2024.03.19.585476

**Authors:** Jasper C. Croll, Hal Caswell

## Abstract

In many species, individuals are embedded in a network of kin with whom they interact. Interactions between kin can affect the survival and fertility rates, and thus the life history of individuals. These interactions indirectly affect both the network of kin and the dynamics of the population. In this way, a non-linear feedback between the kin network and individual vital rates emerges. We describe a framework for integrating these kin interactions into a matrix model by linking the individual kin network to a matrix model. We demonstrate the use of this framework for African elephant populations under varying poaching pressure. For this example, we incorporate effects of the maternal presence and matriarchal age on juvenile survival, and effects of the presence of a sister on young female fecundity. We find that the feedback resulting from the interactions between family members shifts and reduces the expected kin network. The reduction in family size and structure severely reduces the positive effects of family interactions, leading to an additional decrease in population growth rate on top of the direct decrease due to the additional mortality. Our analysis provides a framework that can be applied to a wide range of social species.

## Introduction

From birth to death, almost all organisms are embedded in a network of kin. This kinship network typically includes parents and grandparents at the birth of an individual, and shifts to include children and grandchildren as an individual begins to reproduce. The kinship network extends beyond an individuals’ direct parental line to other related individuals such as sisters, aunts and nieces. The dynamics of the kinship network is closely linked to the life history of individuals and depends on the transition, survival and fertility rates of the individuals within the kinship network (e.g., Caswell, 2019; Caswell and Song, 2021; Coste et al., 2021; Goodman et al., 1974; Jiang et al., 2023).

While all organisms are part of a kinship network, the behavioural and ecological interactions between members of a kinship network are most important in species with a strong social structure (Waldman, 1988). A wide range of plant and animal species exhibit kin interactions, and many of them have the ability to distinguish kin from other individuals. Several evolutionary and ecological mechanisms for the recognition of kin have been proposed, and organisms have been shown to use a wide range of chemical, olfactory, visual and auditory cues to recognize kin (Penn and Frommen, 2010; Tang-Martinez, 2001). Even if individuals are not able to actively recognise kin, regular interactions between kin may occur due to the co-occurrence of kin in space and time. It is debatable whether interactions between kin that arise merely from co-occurrence patterns should be linked to the kin network of an individual or to other spatial and temporal patterns and dynamics of the population (Tang-Martinez, 2001).

Some species actively avoid the co-occurrence of closely related kin to minimize negative effects of kin, such as inbreeding and kin competition (Bengtsson, 1978; Gandon, 1999; Hamilton and May, 1977; Hohenlohe et al., 2021). This avoidance behaviour can result in interesting dispersal patterns, such as sex-specific dispersal distances and behaviour (Li and Kokko, 2019). Conversely, kin of many species cluster together to benefit from positive interactions between family members such as cooperation, alloparenting, knowledge sharing and other forms of assistance (Kramer and Meunier, 2019).

A comprehensive understanding of the genetic diversity and relatedness among individuals is essential to prevent inbreeding and genetic artefacts in a population (Allendorf et al., 2010; Amos and Balmford, 2001; Gobush et al., 2008). Modern techniques using genetic markers provide insight into the current genetic diversity and relatedness of a population. Although these methods can provide an accurate picture of the current genetic diversity of the population, they cannot predict how the relatedness between individuals in the population shift as a result of human and environmental change. The dynamics of the population and the dynamics of the kinship network are strongly linked to changes in relatedness between individuals in the population, and are therefore essential to consider when exploring changes in the relatedness between individuals of a population.

Feedback mechanisms between the life history and the kinship network of an individual can have implications for the conservation and protection of species that live in family groups. Understanding human and environmental influences on the viability of a population is critical for the conservation of a population. The viability of a population is typically assessed by linking individual vital rates to the dynamics of the population through age- or stage-structured models (Akçakaya, 2000; Caswell, 2001). These models are either a linear representation of the system, or only account for interactions at the population level. These interactions are incorporated by making the vital rates dependent on characteristics of the population, such as the population density. This can lead to phenomena such as density dependence, population cycles or Allee effects (Cushing, 2014; Jensen, 1995). However, interactions between individual life histories and other levels of organisation, such as the family group or kinship network, are usually not considered.

Some models incorporate the effect of closely related kin, such as the number of offspring or the presence of the mother as an additional stage structure (Gillespie et al., 2014; Parker et al., 2021; Pavard and Branger, 2012). In these models, stages represent the number of kin of a particular type, and individuals move between the stages as the number of kin of that type changes. Although these models neatly incorporate the direct effect of a specific type of closely related kin on an individual, they do not account for the way these effects cascade through the kin network. For example, if the fecundity of a focal individual depends on the number of sisters she has, then the fecundity of her mother is similarly affected by the number of aunts she has, which in turn affects the number of sisters she has. These cascades through the family are even more complex when life history characteristics are correlated with characteristics of the kin network as a whole, such as the age of the oldest individual in the kin network. In addition, anthropogenic influences can strongly affect the composition of the kin network, thereby altering individual vital rates and the dynamics of the population. These cascades and effects can only be accounted for when the family is modelled explicitly as an additional layer between the individual level and the population level.

In this paper, we present a framework that incorporates feedback mechanisms between the kinship network and the vital rates of an individual. This model links the dynamics of the kinship network to the dynamics and viability of the population. Our model is based on a matrix approach to the formal demography of kinship, which has been extensively developed for human populations (Caswell, 2019, 2020, 2022; Caswell and Song, 2021; Caswell et al., 2023), but the incorporation of kin interactions remains an open problem.

We illustrate the framework by examining how interactions within family groups of African elephants shape the impact of poaching on the viability and relatedness of an elephant population. Female African elephants live in family groups consisting of related kin and are led by a matriarch, who is typically the oldest female in the family group (Athira and Vidya, 2021; Vidya and Sukumar, 2005). The structure of the family groups allows for numerous social interactions between kin, some of which have been well quantified in field studies. Meanwhile, adult females are targeted for ivory poaching, which disrupts the structure of these family groups (Archie and Chiyo, 2012). Thus, poaching has both direct and indirect effects on the viability of the elephant population, resulting from additional mortality and interactions between the family network and individual vital rates. We examine how these interactions shape the response of the population to changes in poaching pressure. Overall, this example not only demonstrates the importance of kin interactions to the conversation for African elephants, but also shows that our framework is a suitable way to incorporate kin interactions into a general matrix approach from conservation.

### A framework to link kin interactions to population dynamics

#### Notation and terminology

Matrices are denoted by uppercase bold characters (e.g., **U**) and vectors by lowercase bold characters (e.g., **x**). Vectors are column vectors by default. The transpose of **x** is denoted as **x**^T^. The vector **1** is a vector of ones. Subscripts are used to refer to a specific entry of a vector; for example, **x**_*i*_ is the *i*th entry of vector **x**. The symbol ◦ denotes the Hadamard, or element-by-element product (implemented by .* in Matlab and by * in R). The function 𝒟(**x**) results in a square matrix with the entries of the vector **x** on the diagonal. The notation ∥**x**∥ denotes the 1-norm of **x**; i.e., the sum of the absolute values of the entries. Occasionally, Matlab notation will be used to refer to rows and columns; for example, **F**(*i*, :) and **F**(:, *j*) refer to the *i*th row and *j*th column of the matrix **F**, respectively.

#### General framework

Demographic models link individual rates (mortality, fertility, growth, development, etc.) to the resulting dynamics at the population level. The kinship network comprises an intermediate level between the individual and the entire population (Fig. 1). Many interactions take place exclusively (or most intensively) among kin. In social species, those interactions can eventually impact the potential dynamics of the population. Interactions among family members, such as alloparenting or food sharing, are incorporated into the model by making the entries in the survival and fertility matrices dependent on the abundance and/or structure of specific types of kin.

**Figure 1:**
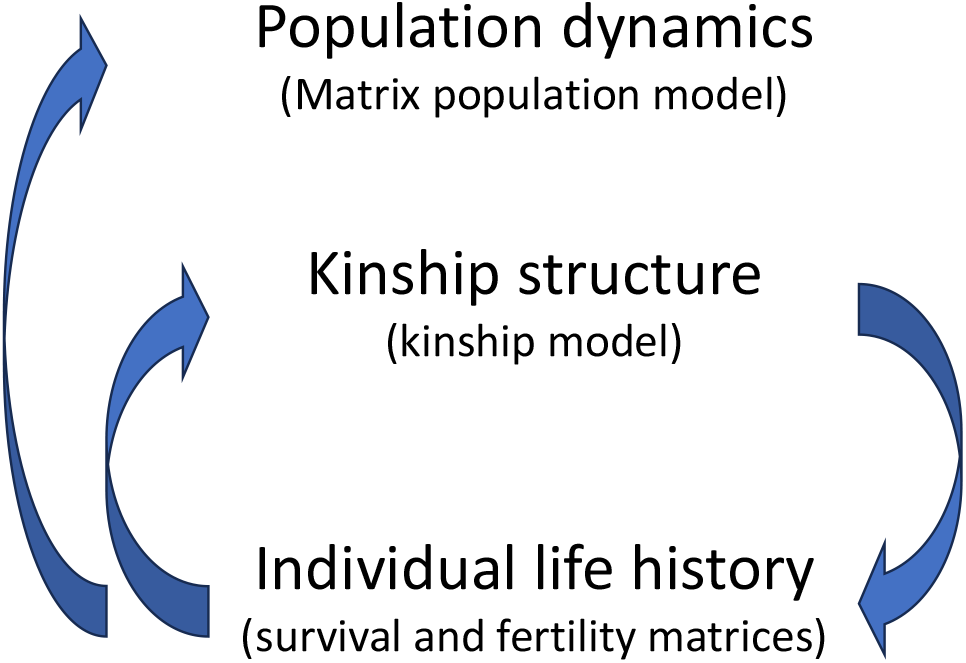
Graphic representation of the interactions between the individual life history, the kinship network and the population dynamics.

In the following sections, we briefly discuss how the population dynamics and kin network are modelled based on the individual transition and survival matrix (**U**) and fertility matrix (**F**). Similar to most matrix model, we focus on projecting the expected structure of the population and kin network and do not project other moments of the distributions. The effect of kin interaction is incorporated in the framework by making the transition and survival matrix and the fertility matrix dependent on the kin network or a specific property hereof. This is a novel way of incorporating family interactions in a matrix model and results in a large set of non-linear equations representing the feedback between individuals and the kin network. In the last section, we describe an iterative approach to solve this set of non-linear equations to obtain an explicit numerical expression of the age-specific kinship network and the corresponding transition and survival matrix and fertility matrix.

#### Population level model

The dynamics of the entire population depend on the transition and survival matrix and the fertility matrix and is given by the familiar population projection matrix **A**:

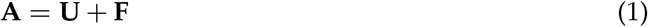

Our framework does not include feedback effects of the population level on the individual vital rates. On the population level, the model therefore behaves as a common linear matrix model. The long-term population growth rate implied by **A** is given by the dominant eigenvalue (λ) of this matrix and is used as a measure of the viability of the population. If λ > 1 the population will grow. If λ < 1 the population will decline to eventual extinction. The eigenvector corresponding to the dominant eigenvalue, normalized to sum to one (**w**), gives the stable age and stage distribution of the population.

#### Kinship level model

The matrix kinship model was introduced in Caswell (2019), and has been extended since then to include a variety of demographic processes, including time variation, multistate life cycles, two-sex models, and losses of kin due to causes of death (Caswell, 2020, 2022; Caswell and Song, 2021; Caswell et al., 2023). The model describes the development of the kinship network of a focal individual, referred to as Focal, as she ages. The age and stage distribution of each type of kin at age *x* of Focal is denoted by a letter, as shown in table 1.

**Table 1:** Recruitment terms and initial age structures of each kin type. The recruitment term for older and younger sisters accounts for the possibility that Focal could be part of a litter. Adapted from Caswell (2019).

| Symbol | Kin type | Initial distribution $\mathbf{k}(0)$ | Subsidy $\beta(x)$ |
| --- | --- | --- | --- |
| **a** | daughters | $\mathbf{0}$ | $\mathbf{F}\phi(x)$ |
| **b** | granddaughters | $\mathbf{0}$ | $\mathbf{F}\mathbf{a}(x)$ |
| **c** | great-granddaughters | $\mathbf{0}$ | $\mathbf{F}\mathbf{b}(x)$ |
| **d** | mothers | $\pi$ | $\mathbf{0}$ |
| **g** | grandmothers | $\sum_i \pi_i \mathbf{d}(i)$ | $\mathbf{0}$ |
| **h** | great-grandmothers | $\sum_i \pi_i \mathbf{g}(i)$ | $\mathbf{0}$ |
| **m** | older sisters | $\sum_i \pi_i \mathbf{a}(i)$ | $\begin{cases} \frac{1}{2} \left[ \frac{\ \mathbf{F}\pi\ }{P(L \geq 1)} - 1 \right] & \text{if } x = 0 \\ \mathbf{0} & \text{if } x > 0 \end{cases}$ |
| **n** | younger sisters | $\mathbf{0}$ | $\begin{cases} \frac{1}{2} \left[ \frac{\ \mathbf{F}\pi\ }{P(L \geq 1)} - 1 \right] & \text{if } x = 0 \\ \mathbf{F}\mathbf{d}(x) & \text{if } x > 0 \end{cases}$ |
| **p** | nieces via older sisters | $\sum_i \pi_i \mathbf{b}(i)$ | $\mathbf{F}\mathbf{m}(x)$ |
| **q** | nieces via younger sisters | $\mathbf{0}$ | $\mathbf{F}\mathbf{n}(x)$ |
| **r** | aunts older than mother | $\sum_i \pi_i \mathbf{m}(i)$ | $\mathbf{0}$ |
| **s** | aunts younger than mother | $\sum_i \pi_i \mathbf{n}(i)$ | $\mathbf{F}\mathbf{g}(x)$ |
| **t** | cousins via older aunts | $\sum_i \pi_i \mathbf{p}(i)$ | $\mathbf{F}\mathbf{r}(x)$ |
| **v** | cousins via younger aunts | $\sum_i \pi_i \mathbf{q}(i)$ | $\mathbf{F}\mathbf{s}(x)$ |

The model projects each type of kin using **U** and **F**. Let **k** denote a generic kin vector, the dynamics are given by

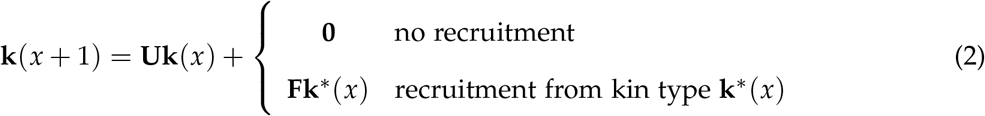

The model is deterministic and provides expected values of the kinship network implied by the vital rates.

The first term in equation (2) describes the survival and ageing of the kin of Focal at age *x*. The second term describes the recruitment of new kin. For some types of kin (e.g., mothers) there is no recruitment of new individuals. For other types, new recruits arrive from the reproduction of some other type of kin (e.g., recruitment of new granddaughters comes from the reproduction of daughters).

We also need the initial age and stage structure of each kin type at the birth of Focal (**k**(0)) (Table 1). Lastly, the vector ***ϕ***(*x*) represents the age and stage structure of Focal, which consists of a one in the row corresponding to the age and stage of Focal (*x*) and zeros elsewhere.

Below, we briefly describe the initial conditions and the recruitment terms for each of the kin considered. The kinship network can be extended as far as desired, but we choose to stop with great-grandparents and great-grandchildren. We give a more detailed description of the kinship model in the online supplementary information, and refer to Caswell (2019) for the derivation of the kinship model.

##### Daughters, granddaughters and great-granddaughters

The initial conditions for daughters, granddaughters, and great-granddaughters are zero. Recruitment of each kin type comes from the reproduction of the previous generation.

##### Mothers, grandmothers and great-grandmothers

The initial condition for mothers is age and stage distribution of mothers at the birth of offspring. There is no recruitment of new mothers. The initial age distributions of Focal’s grandmother and great-grandmother are calculated from the age and stage distribution of Focal’s mother at the birth of Focal. Grandmothers are the mothers of Focal’s mother, and great-grandmothers are the grandmothers of Focal’s mother at that age. There is no recruitment of grandmothers or great-grandmothers.

##### Older and younger sisters

Older sisters are all the daughters of Focal’s mother at Focal’s birth. After Focal’s birth, there can be no further recruitment of older sisters. Younger sisters are daughters of Focal’s mother born after Focal’s birth, so the initial age distribution is zero. Recruitment of younger sisters is the result of the reproduction of Focal’s mother after Focal’s birth. This implies that no sisters are born simultaneously with Focal. This hold for species in which multiple births are rare enough to be ignored.

In species that give birth to multiple offspring in a clutch or litter, Focal may have both older and younger sisters, members of the same clutch, born simultaneously with her. To calculate the expected number of these siblings requires information on, or assumptions about, the frequency distribution of offspring number.

Let *E*(*L*) = ∥**F*π***∥ denote the expected size of a litter. Because we know it must include Focal, Focal’s litter size is conditioned on having at least one member:

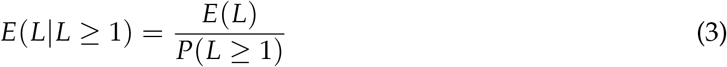

The denominator depends on the distribution of offspring number. For monovular species, *L* is a Bernoulli (0-1) random variable. For species with multiple offspring, the Poisson distribution is an attractive choice. For these cases,

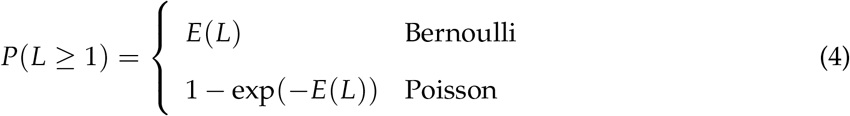

Of course, empirical distributions of litter size can be used if available.

The number of sisters born simultaneously with Focal is the expected clutch size conditioned on the birth of Focal minus one. These sisters are, on average, equally divided between older and younger sisters (Table 1). For monovular species, this reduces to zero, as it should.

##### Nieces, aunts and cousins

The initial distributions of nieces, aunts, and cousins are calculated from the age distribution of Focal’s mother at birth of Focal. Recruitment terms are as given in Table 1.

#### Solving the model

The projected kinship network forms a set denoted with *K*,

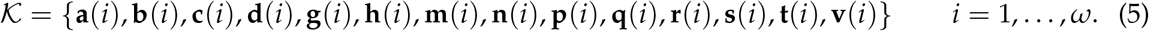

The kinship model described above is a function of the survival and transition matrix (**U**) and the fertility matrix (**F**), producing the set with the kin network.

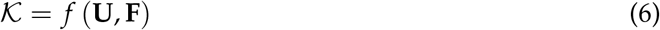

Interaction between family members are incorporated in the framework by making the survival and transition matrix and the fertility matrix dependent on the kin network set or some derived property hereof. This results in a set of non-linear equations, described by the kinship function.

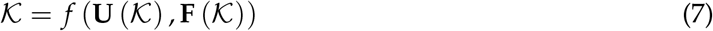

Which we rewrite in the form:

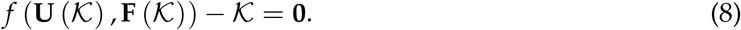

Numerically solving this function will yield the kin network and the corresponding life history matrices. We use the `fsolve` function from the optimization toolbox (The MathWorks Inc., 2022b) in Matlab (The MathWorks Inc., 2022a) to solve equation (8) for 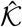. The solution 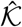 of the kinship network implies a survival and transition matrix **Û** and a fertility matrix 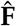. These matrices can be used to evaluate the long term dynamics of the population.

It is tempting to view the solved kin network as a stationary structure. Although the set with the kin structure 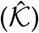 represents the expected kinship structure for all individuals in the population, it is important to keep in mind that the number and structure of a specific type of kin still changes with the age of an individual and therefore with time. The kinship network is therefore only stationary in the sense that the expected distribution of kin for a given age is always the same.

#### Calculating relatedness

Kin types vary in their relatedness to Focal. The kinship model provides the distributions and total count of every kin type at every age of Focal, so we can characterize the kinship network in terms of relatedness. For instance, Focal’s sisters and mother are more closely related to Focal than are Focal’s great-grandmothers or cousins. The described kinship model is based on female relatedness, so it is evident that all sisters in the kinship model share the same mother. In contrast, the model is ambiguous about paternity. This affects the expected relatedness between kin. If two sisters share the same father, their expected relatedness is 0.5 because the sisters overlap in both the maternally and paternally inherited genes. In contrast, if sisters have different and unrelated fathers, the expected relatedness between the sisters drops to 0.25, because they share only maternal genes. For the relatedness to more distant related kin, we assume that fathers are always unrelated. For example, Focal’s sister and Focal might share the same father, but Focal’s nieces are assumed to have a father unrelated to the father of Focal. This results in a minimum and a maximum expected relatedness (ζ^*k*^) between Focal and the kin in the kinship network (Table 2).

**Table 2:** Minimum and maximum expected relatedness between a Focal individual and the various kin types. The maximum expected relatedness assumes that sisters share the same father, while the minimum expected relatedness assumes that all sisters have different, unrelated fathers.

| Symbol | Kin type | maximum relatedness ($\zeta_{max}^k$) | minimum relatedness ($\zeta_{min}^k$) |
| --- | --- | --- | --- |
| **a** | daughters | 0.5 | 0.5 |
| **b** | granddaughters | 0.25 | 0.25 |
| **c** | great-granddaughters | 0.125 | 0.125 |
| **d** | mothers | 0.5 | 0.5 |
| **g** | grandmothers | 0.25 | 0.25 |
| **h** | great-grandmothers | 0.125 | 0.125 |
| **m, n** | sisters | 0.5 | 0.25 |
| **p, q** | nieces | 0.25 | 0.125 |
| **r, s** | aunts | 0.25 | 0.125 |
| **t, v** | cousins | 0.125 | 0.0625 |

The age-specific average relatedness of Focal to the members of the family network (*η*(*x*)) is calculated by weighting the relatedness of a kin type by the relative abundance of the kin type in the family.

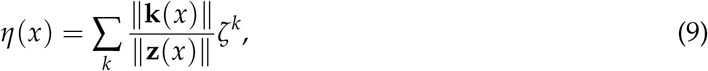

in which **k**(*x*) represents the age distribution of kin of type **k** at age *x* of Focal, ζ^*k*^ represents the relatedness of that kin of type **k** to Focal and **z**(*x*) denotes the age structure of all kin types combined. The expected relatedness of a random individual from the population to its kinship network (*ξ*) is calculated by multiplying the calculated age- and stage-specific average relatedness (*η*(*x*)) with the weighted stable age and stage structure of the population (**w**) to obtain the total expected relatedness of a random individual to the members of their family group:

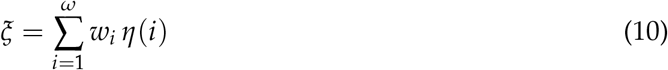

### African elephants: A case study

We demonstrate our framework by exploring how poaching impacts the growth rate of an elephant population while considering several interactions between family members. Female African elephants are known to live in social groups with closely related family members (Athira and Vidya, 2021; Vidya and Sukumar, 2005). Several of these interactions and the effects on the viral rates are well described by observational studies. Meanwhile, the structure of these family groups are disturbed by poaching, targeting females in a specific age range (Wittemyer et al., 2013). Altogether, this makes the application of our framework a valuable exercise, which demonstrate the versatility of our model framework, but also contributes to the insights about secondary effects of poaching on elephant populations.

In the following sections, we describe how we tailored our framework to African elephants. We first outline how we parameterized the model based on measured vital rates of African elephant and how we include poaching as an additional source of mortality. In the following sections, we explain how three different type of family interactions are included in the model. We consider the scenarios in which the presence of a mother influences the juvenile survival (Parker et al., 2021), the presence of sisters increases the fertility of young females Lynch et al. (2019) and matriarch age increases juvenile survival (Foley et al., 2008; Peron et al., 2019). To do so, we use statistical relationships from the literature and derive the required quantities from the kinship model.

#### Vital rates of African elephants

In this study, we parameterise the survival and fertility matrices using age-specific estimates for survival and fertility of female African elephants. We include 63 age classes in our model (*ω* = 63). Survival and fertility values for individuals up to 50 years old are taken from estimates under low poaching conditions by Wittemyer et al. (2021). These estimates are complemented with survival and fertility estimates for females between 51 and 63 years old from the same African elephant population recorded by Wittemyer et al. (2013).

The survival and fertility estimates are smoothed using a Gaussian-weighted moving average with a window of 25 years. After smoothing, the fertility of individuals below 8 years old is set to zero, because females of the African elephant do not mature and reproduce before this age. The resulting survival and fertility matrices are used as a baseline scenario. In this baseline scenario, we treat the impact of poaching as minimal (Supplementary figure S1).

Poaching is included in the model as a proportional decrease in age-specific survival. This corresponds to additive mortality hazard, appropriate for risks like harvesting. To scale poaching mortality, we introduce a parameter *µ* to vary poaching pressure from zero, indicating no poaching, to one, indicating that all individuals in targeted age classes are killed by poaching. In addition, we introduce a vector ***ρ*** of size *ω* describing the relative age-specific vulnerability to poaching, ranging between 0 and 1. Poaching pressure for juveniles below 8 years old is much lower than poaching pressure of older females. The poaching vulnerability of females below 8 years old is therefore set to zero (***ρ***(1 : 8) = 0). Similarly, adolescents between 9 and 18 years old are approximately half as likely to die from poaching than older females. The poaching vulnerability of females between 9 and 18 years old is therefore set to 0.5 (***ρ***(9 : 18) = 0.5). The poaching vulnerability for females above 18 years old is set to one (***ρ***(19 : 63) = 1) (Wittemyer et al., 2013). The survival matrix including poaching (**U**^*µ*^) is given by

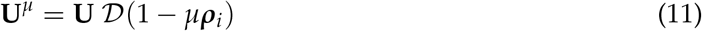

We will indicate a specific poaching pressure by adding a superscript to matrices and vectors (e.g. **U**^*µ*^, **k**^*µ*^).

In the model analysis, we solve the age-specific kinship network and calculate the population growth rate under varying poaching conditions, ranging from no poaching to high poaching (*µ* = 0 to *µ* = 0.2, with steps of 0.002) for each scenario. Additionally, we calculate the expected relatedness between an individual and their kin network, to gain insight in the effect of poaching on the kin network of an individual.

#### Effect of mother presence on juvenile survival

We first consider the effect of the presence of a mother on the survival of juveniles (Parker et al., 2021). Because individuals have exactly one mother, the presence of the mother follows a binomial distribution. The probability of having a living mother at age *x* therefore directly follows from the expected number of mothers projected by the kinship model:

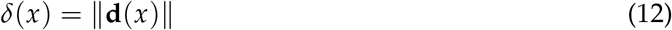

We use the statistical relation between mother presence and survival quantified by Parker et al. (2021) to construct a vector ***α*** representing the age-specific proportional decrease in survival due to the absence of a mother (Supplementary table S1). Juvenile elephants up to two years old do not survive in the absence of their mother (***α***(1 : 2) = 1). The survival of juveniles between two and eight years old is reduced by 14.27 percent (***α***(3 : 8) = 0.1427). The survival of females between nine and eighteen years old decreases by 3.87 percent (***α***(9 : 18) = 0.0387) in the absence of their mother. The presence of the mother does not affect the survival of older females (***α***(19 : 63) = 0).

We use the probability of having a living mother under a specific poaching pressure *δ*^*µ*^(*x*) relative to the probability of having a living mother without poaching (*δ*^0^(*x*)). This guarantees that the survival matrix without poaching is equal to the baseline survival matrix derived from the observational data 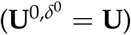, resulting in the follow equation for the survival matrix.

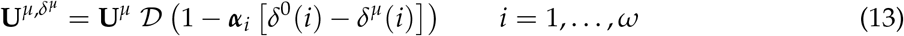

#### Effect of sister presence on fertility

The second interaction we consider is the effect of the presence of a sister on the fecundity of young females (Lynch et al., 2019). The probability distribution of the number of sisters at a given age (*υ*(*x*)) is required to derive the probability that an individual has at least one sister. The kinship model only projects the expected number and age distribution of sisters. For humans, the probability distribution of the number of sisters is closely approximated by a Poisson distribution (Caswell, 2024). Because the vital rates of elephants are very similar to those of humans, we also use a Poisson distribution to approximate the probability distribution of the number of sisters. From this, we can derive an expression for the probability an individual of age *x* has at least one sister:

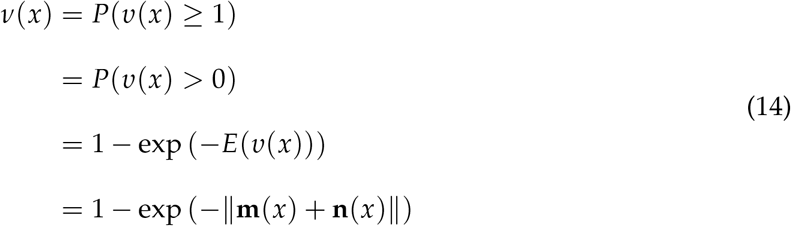

We use the statistical relationship between the presence of a sister and the fertility of an individual, quantified by Lynch et al. (2019), to construct a vector ***β*** representing the age-specific fractional increase in fertility due to the presence of at least one sister (supplementary table S2). The values in this vector are calculated as the difference in the probability of successfully reproducing with and without at least one sister around, divided by the probability of successfully reproducing without a sister around.

We use the probability of having at least one sister under a specific poaching pressure (*ν*^*µ*^(*x*)) relative to the probability of having at least one sister without poaching (*δ*^0^(*x*)). This guarantees that the fertility matrix without poaching is equal to the baseline fertility matrix derived from the observational data 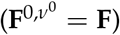, resulting in the follow equation for the survival matrix.

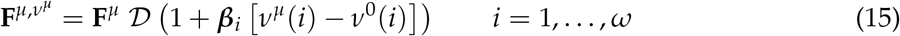

#### Effect of matriarch age on juvenile survival

The third interaction we consider is the effect of the oldest age in the family on juvenile survival (Foley et al., 2008; Peron et al., 2019). We can derive the expected oldest age in the family 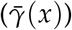 from the expected age distribution using order statistics (Wilks, 1948). The expected age distribution of the entire kin network (**z**(*x*)) is calculated by summing the age distributions of all kin projected by the kinship model.

Because we conditioned the model on the survival of Focal, we know that the oldest age in the family is at least as old as the age of Focal (i.e., *γ*(*x*) ≥ *x*). The probability that the oldest age in the family (*γ*(*x*)) is equal or below a certain age *y* given the age of Focal (*x*) is the same as the probability that all individuals older than Focal have an age below *y*:

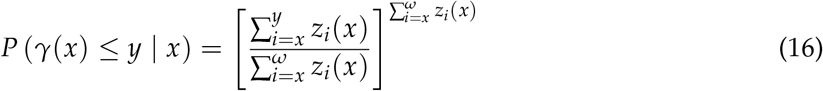

The denominator and the power in this formula are the total number of individuals with an age equal to or older than the age of Focal. The numerator is the total number of individuals with an age below or equal to *y* and above or equal to the age of Focal (*x*).

The probability that the oldest age in the family is exactly equal to a specific age *y* is the probability that the oldest age is equal or below *y* minus the probability that the age is below *y*:

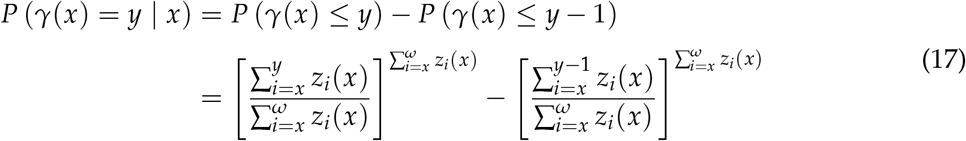

It is important to note that these calculations are only relevant for ages above Focal’s age (*y* > *x*) as individuals younger than Focal cannot be the oldest individual in the family (*P*(*γ*(*x*) = *y* | *y* < *x*) = 0). From the probability distribution of the oldest age in the family we can calculate the expected oldest age in the family, given a specific age of Focal:

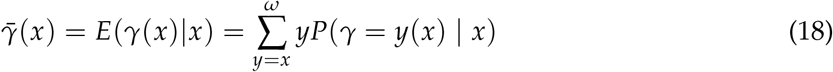

We use the logit-relationship statically derived by Peron et al. (2019) to link the expected oldest age in the family to the proportional increase in survival due to the oldest age in the family (*θ*). We use the expected oldest age in the family under a specific poaching pressure 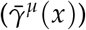 relative to the expected oldest age in the family without poaching 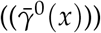. This guarantees that the fractional increase without poaching is always equal to one and the survival matrix without poaching is equal to the baseline survival matrix derived from the observational data 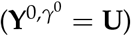.

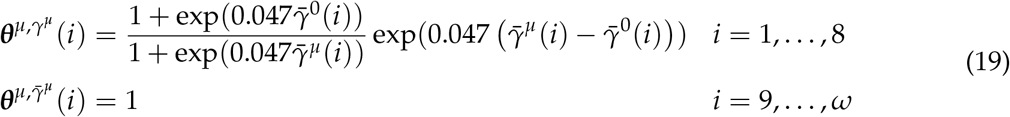

The survival matrix is then given by

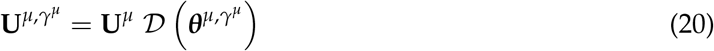

### Model results for African elephants

We introduced a novel way to incorporate family interactions into matrix models for conservation. As an example, we tailored the model to study the effect of poaching on the kin network and population dynamics while accounting for three different family interactions. Here we discuss the results from our model application to African elephants. We first discuss the population level result and afterwards discuss the specific mechanisms acting on the family level.

#### Population growth effects

Poaching reduces the growth rate of the elephant population (Fig. 2). In the absence of family feedback, the population will grow as long as the poaching pressure is below approximately 0.095, and will decrease if poaching pressure is above this value. The feedbacks between kinship structure and the individual life histories all amplify the effect of poaching on population growth. This amplification occurs because poaching alters the family structure, which diminishes the positive effects from the family interactions. As a consequence, poaching has a stronger negative effect on the population growth when these interactions are considered. Family feedback mechanisms reduce the critical poaching pressure above which the elephant population will decrease in size (Fig. 2).

**Figure 2:**
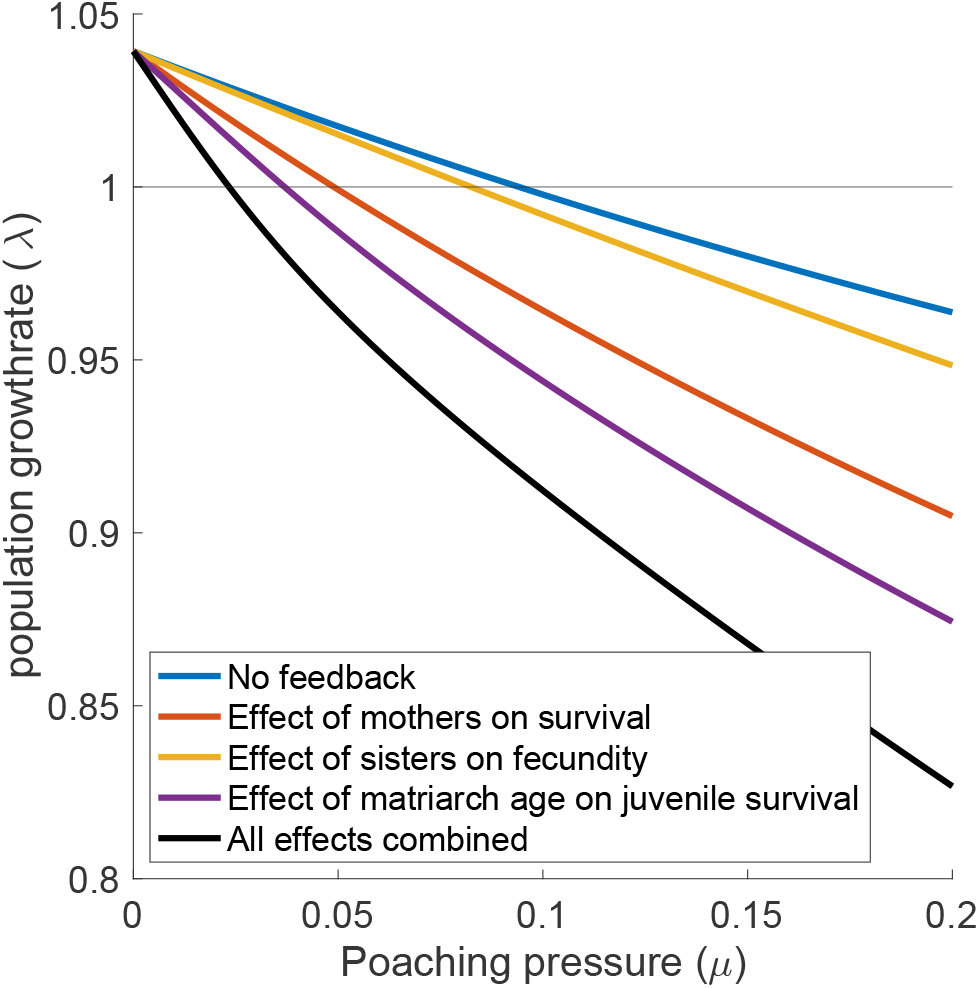
Growth rate of the population under increasing poaching pressure with various interactions between family members.

#### Relatedness

As Focal ages, her average relatedness to her family group exhibits three distinct stages, which are mainly driven by the birth and death of kin in the maternal line of Focal (e.g. mothers, grandmothers, daughters and granddaughters) (Fig. 3). Initially, the average relatedness decreases with the age of Focal during the juvenile period. This pattern arises because some of the most closely related kin (e.g., mothers and grandmothers) of Focal might die, while Focal produces no new closely related kin in the form of daughters. The average relatedness to the kinship network begins to increase as soon as Focal starts to reproduce, because the daughters of Focal are always closely related to her. At old age, the average relatedness to the kinship network decreases again, because the death of Focal’s daughters leaves only more distant relatives such as great-granddaughters and nieces.

**Figure 3:**
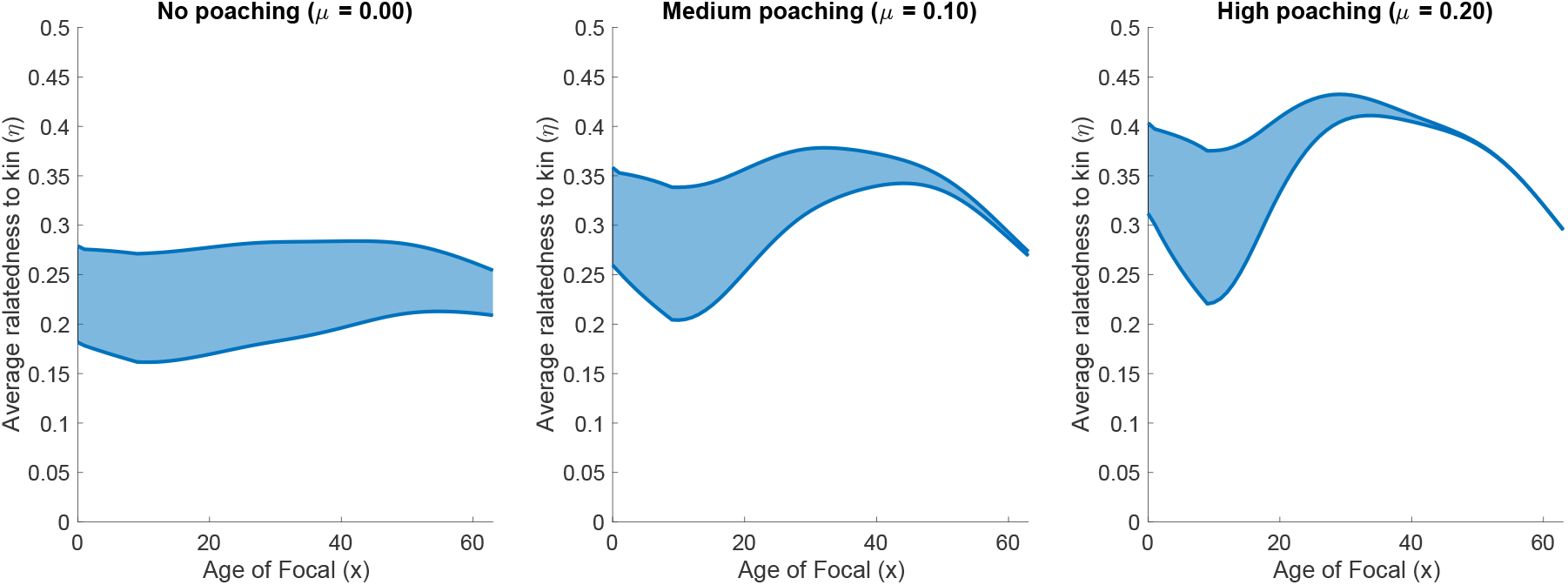
Average relatedness to the kinship network throughout the life of a Focal individual under three different poaching intensities. The lowest relatedness values are calculated with the assumption that all fathers are unrelated, while the highest relatedness values are calculated with the assumption that all offspring of an individual have the same father.

An increase in poaching pressure increases the average relatedness to the kinship network, and also amplifies the increase and decrease in relatedness throughout the life of Focal. These patterns emerge because poaching decreases the number of distantly related individuals such as aunts, cousins and nieces to a disproportionately large extent compared to kin in the maternal line of Focal such as mothers and daughters.

Figure 4 shows the relationship between poaching pressure and the average relatedness to the family group, as affected by the various family feedback effects. The expected relatedness increases with poaching pressure. The family feedbacks all increase the expected relatedness of Focal to her kinship network, because all these interactions amplify the effect of poaching.

**Figure 4:**
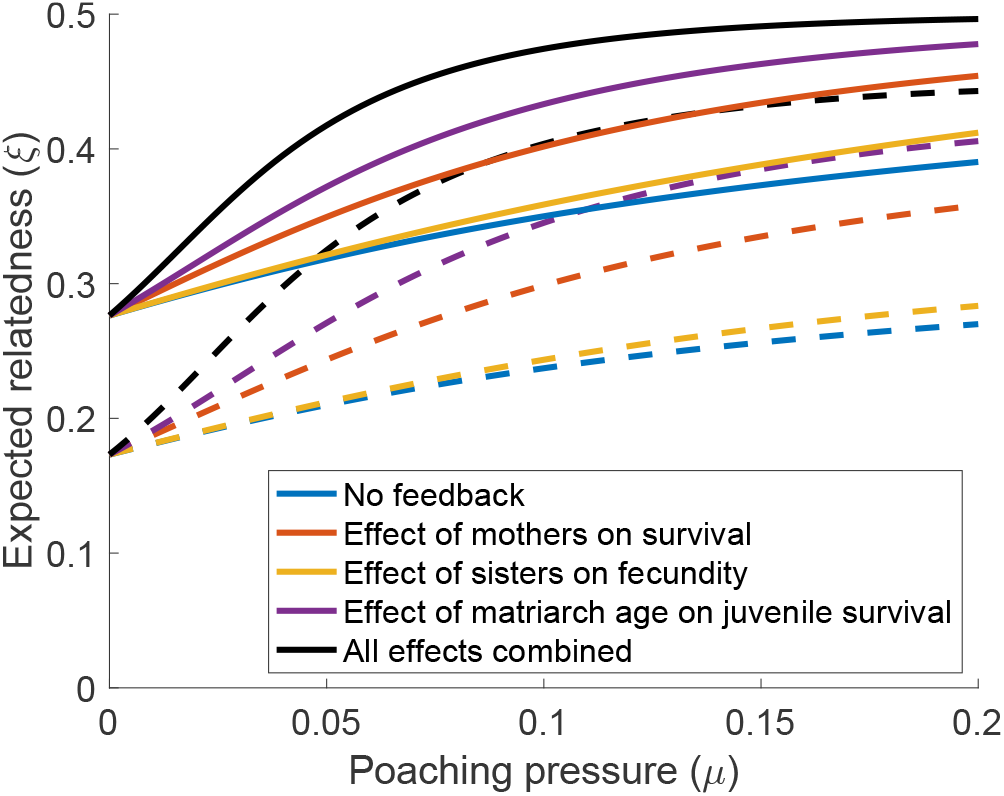
Expected relatedness of an individual in the population to the family group of that individual, computed with various types of feedback mechanisms. Solid lines are calculated with a high relatedness in which sisters all have the same father, and dashed lines are calculated with a low relatedness in which all fathers are completely unrelated.

#### Effect of mother presence on juvenile survival

The presence of a mother increases the survival of juvenile individuals. Poaching strongly decreases the presence of the mother of Focal (Fig. 5). The feedback between the survival of Focal and the presence of her mother does not strongly affect the presence of the mother itself (Fig. 5) or other direct ancestors such as grandmothers and great-grandmothers. However, the additional feedback of mothers on juvenile survival substantially decreases the number of other kin such as daughters, granddaughters, sisters, aunts and nieces (Supplementary figure S2). Consequently, the feedback of mothers on the survival of juveniles amplifies the impact of poaching on the population growth rate (Fig. 2).

**Figure 5:**
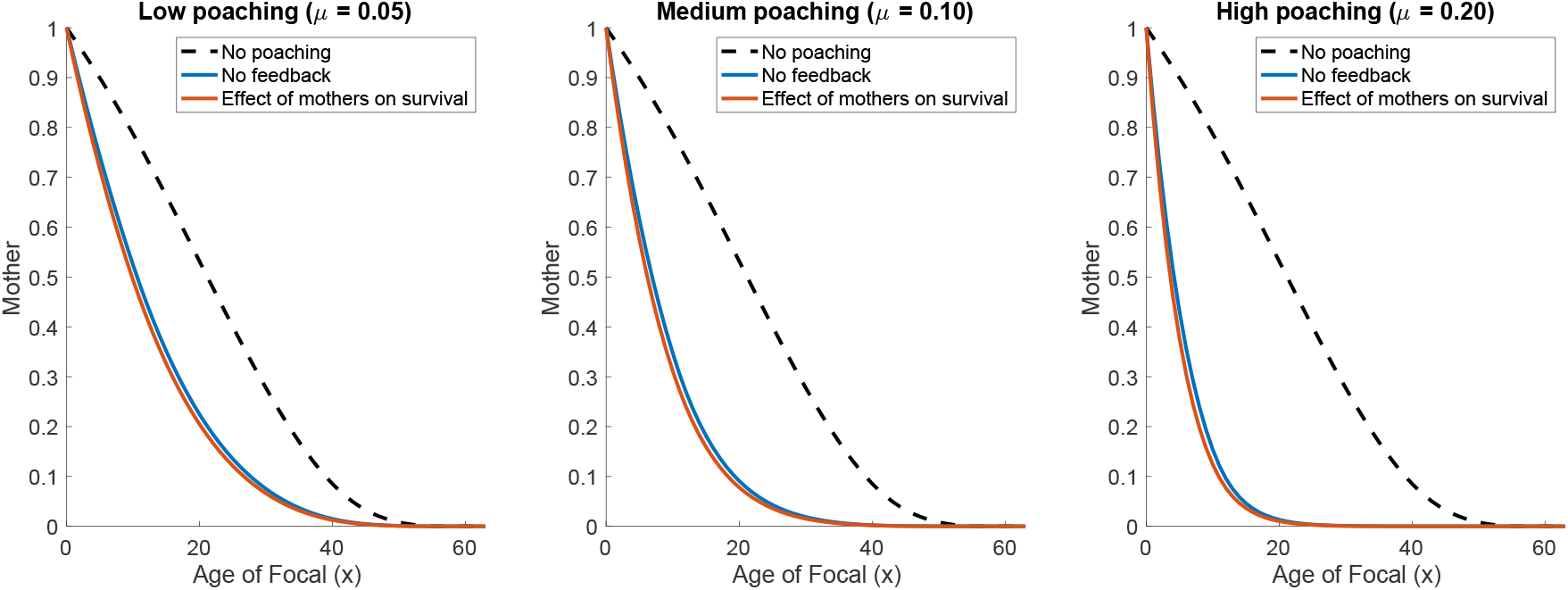
Expected presence of Focal’s mother throughout her life, under various poaching pressures and with and without the feedback of mothers on the survival of juveniles. The dynamics of the extended kinship network throughout the life of Focal is shown in supplementary figure S2.

#### Effect of sister presence on fertility

The fertility of individuals up to 20 years of age decreases if they have fewer than one sister around. At young age, the expected number of sisters, and thus the probability that Focal has at least one sister, increases with age, because Focal’s mother produces new sisters. At older ages, the expected number of sisters, and thus the probability of having at least one sister, declines because older sisters die off, and no new sisters are produced any more (Fig. 6). Poaching strongly decreases the number of sisters of Focal, and thus the probability that Focal has at least one sister. The feedback between family structure and the survival slightly decreases the probability that at least one sister is present a bit further, but also affects the expected number of other family members (Supplementary figure S3). As a result, the effect of the presence of a sister on fertility amplifies the effect of poaching on the population growth rate (Fig. 2).

**Figure 6:**
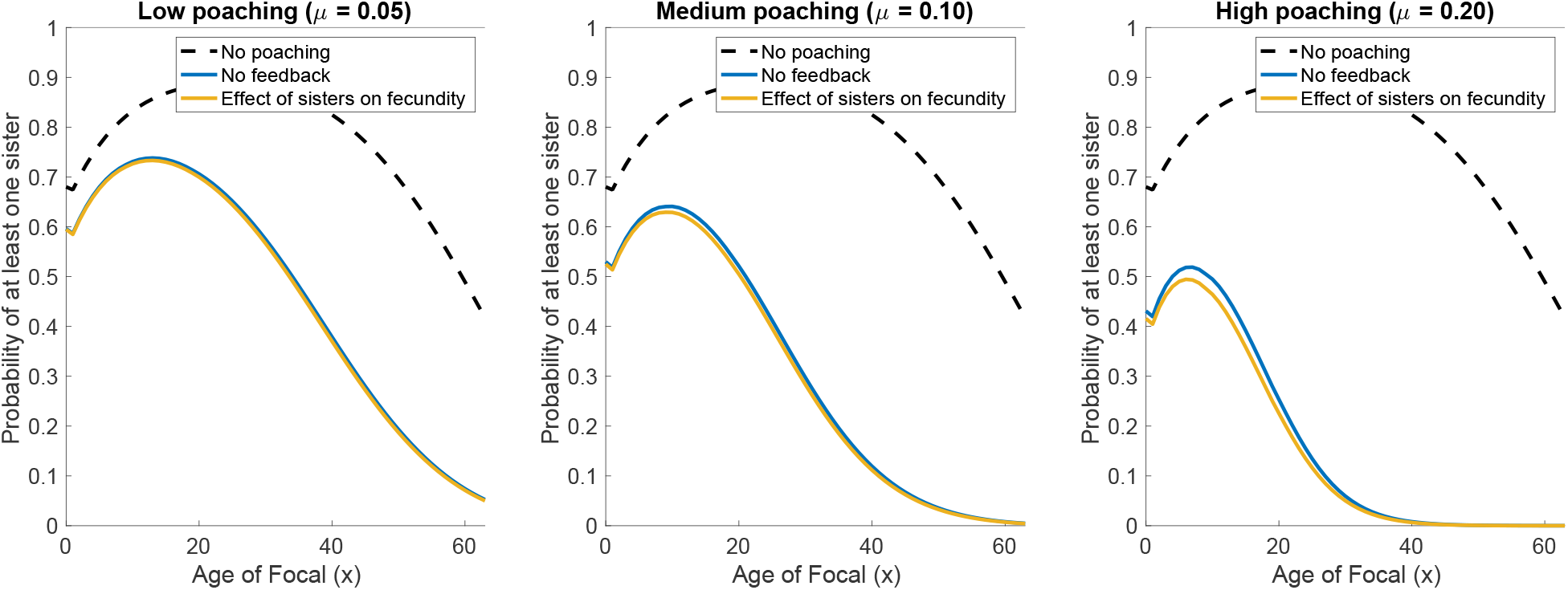
Probability of the presence of at least one sister as a function of the age of Focal, for various poaching pressures and with and without the feedback of sisters on fertility. The dynamics of the extended kinship network throughout the life of Focal is shown in supplementary figure S3.

#### Effect of matriarch age on juvenile survival

The survival of juveniles increases with the oldest age in the family. As Focal grows older, the oldest age in the family increases and converges to the age of Focal (Fig. 7). This pattern occurs because the analysis focuses on the kin of Focal, and thus is conditioned on the survival of Focal. The older Focal is, the more likely it is that Focal herself is the oldest female in the kinship network and therefore the matriarch.

**Figure 7:**
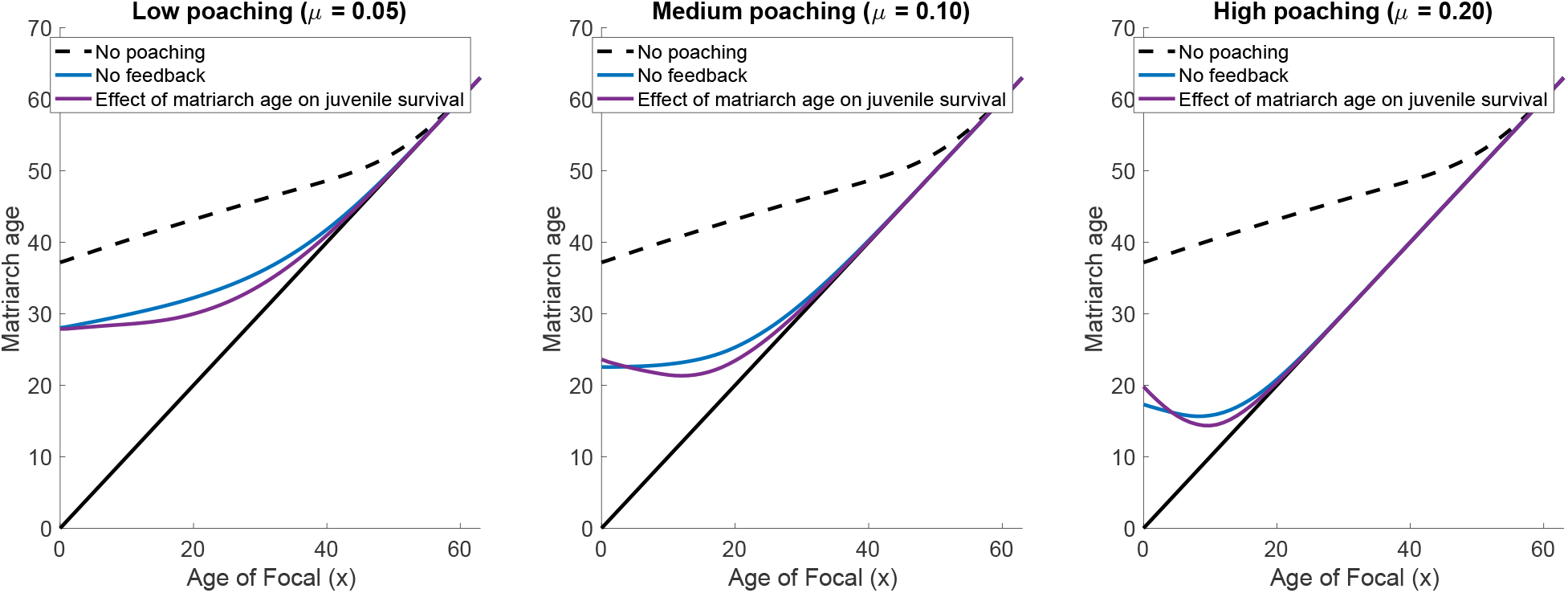
Expected oldest age in the kinship network, as a function of the age of Focal for various poaching pressures and with and without the feedback of mothers on the survival of juveniles. The complete kinship network throughout the life of Focal is shown in supplementary figure S4.

Poaching reduces the number and the life expectancy of kin, resulting in a decrease in the expected oldest age in the kinship network. Additionally, poaching increases the likelihood that Focal’s mother is the matriarch of the family. Meanwhile, poaching also reduces the survival of Focal’s mother, possibly causing an additional decrease in the expected oldest age in the kinship network during the first years of Focal’s life (Fig. 7).

The age of the matriarch affects the survival of juveniles. Incorporating this feedback further reduces the number of older kin, such as older sisters and aunts, which causes the expected oldest age to drop further during Focal’s life (Supplementary figure S4). Interestingly, the feedback also shifts the age distribution of Focal’s mother at the birth of Focal towards older ages, which counteracts the effect of the low number of kin and slightly increases the expected oldest age in the family at the birth of Focal. Regardless of these small particular effects, the effect of the age of the matriarch on juvenile survival reduces population growth rate, as a function of poaching pressure, more than any of the other feedbacks (Fig. 2).

## Discussion

Interactions between kin, such as cooperation, alloparenting, knowledge sharing and other forms of assistance, can affect the vital rates of individuals (e.g. Bengtsson, 1978; Hamilton and May, 1977; Kramer and Meunier, 2019; Waldman, 1988). Since the dynamics of a population reflect the life histories of individuals, we can intuitively understand that the structure of a family network could strongly influence the viability and dynamics of a population. Meanwhile, the structure of a family network is also shaped by the life histories of the family members (Caswell, 2019), creating opportunities for feedback mechanisms between individual life histories, family structures, and population dynamics. These feedback mechanisms may influence how the viability and dynamics of populations with strong family interactions will respond to anthropogenic influences. Here, we provide a framework for modelling populations with strong kin interactions by combining the matrix model for a kinship network (Caswell, 2019) with a matrix population model (Caswell, 2001) based on the same vital rates. The kinship model utilizes the vital rates from the population model to describe the abundance and dynamics of kin throughout the life of a focal individual. Meanwhile, the vital rates in the population model depend on specific parts of the kinship network calculated with the kinship model. Because of the interdependence between the kinship model and the population model, the kinship network and population structure can only be obtained numerically. It is important to note that the obtained family structure of individuals is not stationary in the same way as the structure of the population. The population structure gives the proportion of individuals at each age and stage and is time invariant. The solved kinship network gives the expected number and structure of each kin type at a given age of Focal. The number and structure of kin changes with the age of Focal, and therefore changes with time. However, the solved kin network is constant in the sense that all individuals of the same age have the same expected kin network. This is sufficient to derive population-level quantities such as the long-term population growth rate, which is a common measure for the viability of the population, and the average relatedness between individuals.

Our model deals with the long term expected kin network, relatedness and population growth rate, which is the most common way to use matrix population models in ecology and conservation (Caswell, 2001). As a result, the expected age-specific kin network of all individuals in the model is the same. In real populations, the number and structure of kin might differ between individuals. On the population level, this would result in a probability distribution of the number of kin. For example, we might exactly know the number of sisters of a specific individual from observations, but the number of sisters of all individuals in the population with a specific age will have some probability distribution. The shapes of these probability distributions are usually unknown, and our model deals with the expected value of the probability distributions. We did require additional information about the probability distribution of sisters in the population to calculate the probability that an individual has at least one sister. For humans, the distribution of sisters is commonly approximated with a Poisson distribution (Caswell, 2024; Feng et al., 2023; Song et al., 2015; Song and Mare, 2019). Because the vital rates of elephants are close to the vital rates of humans, we used a Poisson distribution for the number of sisters in our example of African elephants. It requires extensive field studies to demonstrate whether this holds for other animals as well. Without interactions, it is possible to model the dynamics of other moments of the probability distributions of the kin network, such as the variances (Caswell, 2024; Pollard, 1966). Addition of family interactions to the stochastic kinship model to predict other moments of the probability distributions requires numerous linear approximations of the kin interactions and remains an open question.

Our analysis is based on the one-sex version of the kinship model, as is appropriate for a matriarchically-organized species like the African elephant. However, multi-state and two-sex versions of the model (Caswell, 2020, 2022) make it possible to apply our modelling framework to species with a wide range of population structures and family interactions.

Our analysis of African elephants provides an example of how the model provides insight in the effect of individual interactions on population growth. In our model, we focused in particular on the response of the family and population to poaching. The analysis is possible because of extensive field studies documenting the vital rates and interactions among female African elephants: an increase in juvenile survival due to the presence of their mother (Parker et al., 2021), an increase in fertility of young females due to the presence of a sister (Lynch et al., 2019), and an increase in juvenile survival with the age of the oldest female in the family group (Foley et al., 2008; Peron et al., 2019).

These positive interactions between family members amplify the negative effect of poaching on the population growth rate. It is no surprise that poaching, as an additional mortality source, reduces the population growth rate. However, poaching also damages the structure of the kinship network (an effect commonly observed in wild populations of elephants (Aleper and Moe, 2006; Barnes and Kapela, 1991; Foley, 2002; Mkuburo et al., 2020)). The result is a decrease in the survival and fertility of individuals in the population, which adds to the direct effect of poaching mortality. This effect is often invoked for mortality effects in social species; Figure 2 shows a quantitative example.

Our analysis does not include the potentially important processes of fission and fusion of elephant family groups (Archie et al., 2011; Wittemyer et al., 2005). Fission and fusion might especially affect the oldest age in the family and the relatedness between individuals. Although core family units of African elephants are strong and their social networks robust (Goldenberg et al., 2016), family groups might blend temporarily based on environmental conditions and group size (Wittemyer et al., 2005). Large family groups tend to split based on the relatedness, with closely related individuals more likely to end up in the same group (Archie et al., 2011; Wittemyer et al., 2009). This process is roughly captured by limiting the family network model to Focal’s great-grandmother, Focal’s grandmother, and the descendants of Focal’s grandmother. The absence of more distantly related family members could reflect a split of the family network based on relatedness. On the other hand, our model does not capture the fusion of small family groups. Although more closely related family groups are more likely to fuse (Archie et al., 2011; Wittemyer et al., 2009), the relatedness between these individuals might still be relatively low. Incorporating fission and fusion of family groups into the matrix kinship model is an open research problem.

Genetic relatedness between individuals can play a crucial role in the conservation of wildlife populations (Hohenlohe et al., 2021). The matrix kinship model offers the opportunity to predict relatedness within family groups based on individual vital rates. As such, it could serve as a null model for studies of relatedness within populations. The pairwise relatedness between individuals predicted by our model for African elephants is very similar to the average pairwise relatedness within observed family groups (Archie et al., 2006; Wittemyer et al., 2009).

Two other examples of the amplification of demographic perturbations by kin interactions give an idea of how general the effect may be. Verdery et al. (2020) derived a ‘bereavement multiplier’ for the effect of deaths due to COVID-19 in the United States. Using a kinship model, they found that for every COVID death, approximately 9 bereaved individuals will have lost a grandparent, parent, sibling, spouse, or child. Such losses are known to have a variety of damaging effect on the bereaved (e.g., review in Caswell et al., 2023). In a quite different species, the Wandering Albatross forms long-lasting pair bonds, which can be broken by mortality of one partner (females are subject to higher mortality due to fishery bycatch) or by divorce. Of course, bycatch mortality reduces the population growth rate. However, Sun et al. (2022) found that this effect was amplified by the effect on the surviving partner; with large reductions in lifetime reproductive success of the male survivors of pair bond disruption. Amplification of mortality factors by connections within a kinship network is probably more relevant than is recognized.

Accounting for the effects of family interactions on population growth of social species requires a level between that of the individual and that of the population. The framework presented here incorporates family interactions in the dynamics of populations while using the same tools used for population projections. The framework can provide new insight and hypothesis about the importance of interactions between family interactions and is a first step towards the inclusion of family interactions in projection models for ecological predictions and management of populations.

## Supporting information

Matlab code to run the model

## Acknowledgments

This research was supported by the European Research Council under the European Union’s Horizon 2020

## Statement of Authorship

JCC conceived the initial concept of this study, analysed the model and wrote the original draft. Both JCC and HC contributed to the model formulation and contributed to the writing of all stages of the manuscript.

## Data and Code Availability

This paper does not include new data. All Matlab code needed to reproduce the results are included as supplementary material with this publication.

## S1 Elaborate description of the kinship model

In this section, we give a more detailed explanation of the equations in table 1 from the main text. The derivations of the equations are explained entirely in Caswell (2019).

### Daughters, granddaughters and great-granddaughters

Daughters (**a**(*x*)), granddaughters (**b**(*x*)) and great-granddaughters (**c**(*x*)) of Focal are only born after Focal. The initial distribution of these kin types at the birth of Focal is therefore always zero (**k**(0) = 0). The subsidy term of these kin types consists of the distribution of the parent kin multiplied by the fertility matrix. For example, in the case of Focal’s daughters, the subsidy term consists of the distribution of Focal multiplied by the fertility matrix (***β***(*x*) = **F*ϕ***(*x*)). Similarly, the subsidy term for Focal’s granddaughters is the distribution of daughters multiplied by the fertility matrix (***β***(*x*) = **Fa**(*x*)) and the subsidy term for Focal’s great-granddaughters is the distribution of granddaughters multiplied by the fertility matrix (***β***(*x*) = **Fb**(*x*)).

### Mothers, grandmothers and great-grandmothers

The expected age structure of mothers (**d**(*x*)) at the birth of Focal (***π***) is the distribution of mothers at the birth of children. It is calculated by weighting the stable structure of the population (**w**) by the fertility and normalizing the resulting vector to sum to one.

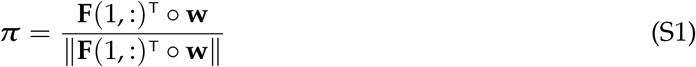

As Focal does not acquire additional mothers during its life, the subsidy term of Focal’s mother is zero (***β***(*x*) = 0). The distribution of mothers throughout Focal’s life is calculated using only the survival matrix.

Focal’s grandmother (**g**(*x*)) is the mother of Focal’s mother. Assuming equal family distributions for all individuals, the initial distribution of Focal’s grandmother is calculated by weighting the distribution of the mother by the initial distribution of mothers at the birth of Focal (**g**(0) = ∑_*i*_ ***π***_*i*_**d**(*i*)).

Similarly, the great-grandmother (**h**(*x*)) of Focal is the grandmother of Focal’s mother. The initial distribution of great-grandmothers is therefore calculated by weighting the distribution of grandmothers with the initial distribution of mothers (**h**(0) = ∑_*i*_ ***π***_*i*_**g**(*i*)). Focal does not acquire new grandmothers or great-grandmothers throughout its life, and the subsidy term of these kin is therefore zero (***β***(*x*) = 0).

### Sisters

Sisters older than Focal (**m**(*x*)) are the daughters of Focal’s mother, born before Focal’s birth. The initial distribution of older sisters is therefore calculated as the distribution of daughters weighted by the distribution of mothers at Focal’s birth (**m**(0) = ∑_*i*_ ***π***_*i*_**a**(*i*)). After Focal’s birth, no new older sisters are born, resulting in a subsidy term of zero for older sisters (***β***(*x*) = 0).

Younger sisters (**n**(*x*)) are born after Focal’s birth, and the initial distribution of younger sisters is therefore zero (**n**(0) = 0). Younger sisters are the offspring of Focal’s mother born after Focal, and the subsidy term for younger sisters is therefore the age distribution of mothers multiplied by the fertility matrix (***β***(*x*) = **Fd**(*x*)).

The offspring of Focal’s mother born in the first time step (*x* = 0) include Focal, but might also include sisters of Focal if Focal’s mother gives birth to multiple offspring at the same time. The expected clutch size (*E*(*L*)) with Focal is the expected number of offspring conditioned on the birth of Focal (*E*(*L*|*L* ≥ 1)). This is calculated by correcting the expected clutch size for the probability a clutch is born (*P*(*L* > 0)). In this case, we can approximate the probability distribution of the number of offspring with a Poisson distribution:

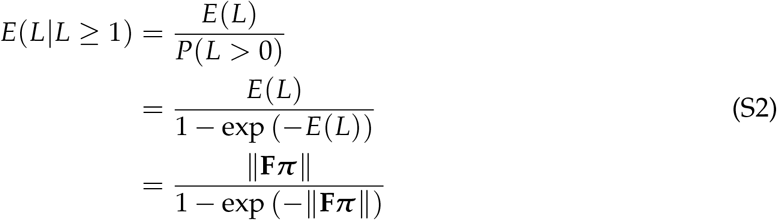

The number of sisters born simultaneously with Focal is the expected clutch size conditioned on the birth of Focal minus one. These sisters are, on average, equally divided between older and younger sisters.

### Nieces, aunts and cousins

The nieces of Focal are the offspring of Focal’s older and younger sisters (**p**(*x*) and **q**(*x*) respectively). Focal’s older sisters could produce offspring before Focal’s birth. The nieces at Focal’s birth are the granddaughters of Focal’s mother at the birth of Focal. We calculate the initial distribution of nieces through older sisters by weighting the distribution of granddaughters with the distribution of mothers at the birth of Focal (**p**(0) = ∑_*i*_ ***π***_*i*_**b**(*i*)). Older sisters might still produce offspring after Focal’s birth, and the subsidy term of these nieces is therefore the distribution of older sisters multiplied by the fertility matrix (***β***(*x*) = **Fm**(*x*)).

Younger sisters are not yet born at the time of Focal’s birth, so the initial distribution of nieces through younger sisters is zero (**p**(0) = 0). The subsidy term of nieces through younger sisters is the distribution of younger sisters multiplied by the fertility matrix (***β***(*x*) = **Fn**(*x*)).

The aunts of Focal are divided into aunts older than Focal’s mother (**r**(*x*)) and aunts younger than Focal’s mother (**s**(*x*)). Aunts older than Focal’s mother are the older sisters of Focal’s mother. We calculate the initial distribution of these aunts by weighting the distribution of older sisters with the distribution of mothers at the birth of Focal (**r**(0) = ∑_*i*_ ***π***_*i*_**m**(*i*)). Aunts older than Focal’s mother cannot be born after Focal’s birth, so the subsidy term for these aunts is zero (***β***(*x*) = 0). Aunts younger than Focal’s mother are the younger sisters of Focal’s mother. The initial distribution of these aunts is therefore calculated by weighting the distribution of younger sisters with the distribution of mothers at the birth of Focal (**s**(0) = ∑_*i*_ ***π***_*i*_**n**(*i*)). Aunts younger than Focal’s mother can still be born after Focal’s birth. The subsidy term of these aunts is therefore the distribution of grandmothers multiplied by the fertility matrix (***β***(*x*) = **Fg**(*x*)).

Lastly, both the aunts older than Focal’s mother and aunts younger than Focal’s mother produce cousins of Focal (**t**(*x*) and **v**(*x*) respectively). The cousins of Focal are the nieces from Focal’s mother. The initial distribution of these cousins is calculated by weighting the distribution of nieces with the distribution of mothers at the birth of Focal (**t**(0) = ∑_*i*_ ***π***_*i*_**p**(*i*) and **v**(0) =∑_*i*_ ***π***_*i*_**q**(*i*)). The subsidy terms for cousins are the distribution of aunts older than Focal’s mother multiplied by the fertility matrix (***β***(*x*) = **Fr**(*x*)) and the distribution of aunts younger than Focal’s mother multiplied by the fertility matrix (***β***(*x*) = **Fs**(*x*)).

## S2 Supplementary tables and figures

**Table S1:** Calculation of the proportional decrease in survival (***α***) due to the absence of the mother based on mean survival values estimated by Parker et al. (2021).

| Age group | Estimated mean<br>survival with<br>mother | Estimated mean<br>survival without<br>mother | Proportional<br>decrease used in<br>our model ($\alpha$) |
| --- | --- | --- | --- |
| 0-2 | 0.970 | 0.000 | 1.000 |
| 3-8 | 0.988 | 0.847 | 0.1427 |
| 9-18 | 0.981 | 0.943 | 0.0387 |

**Figure S1:**
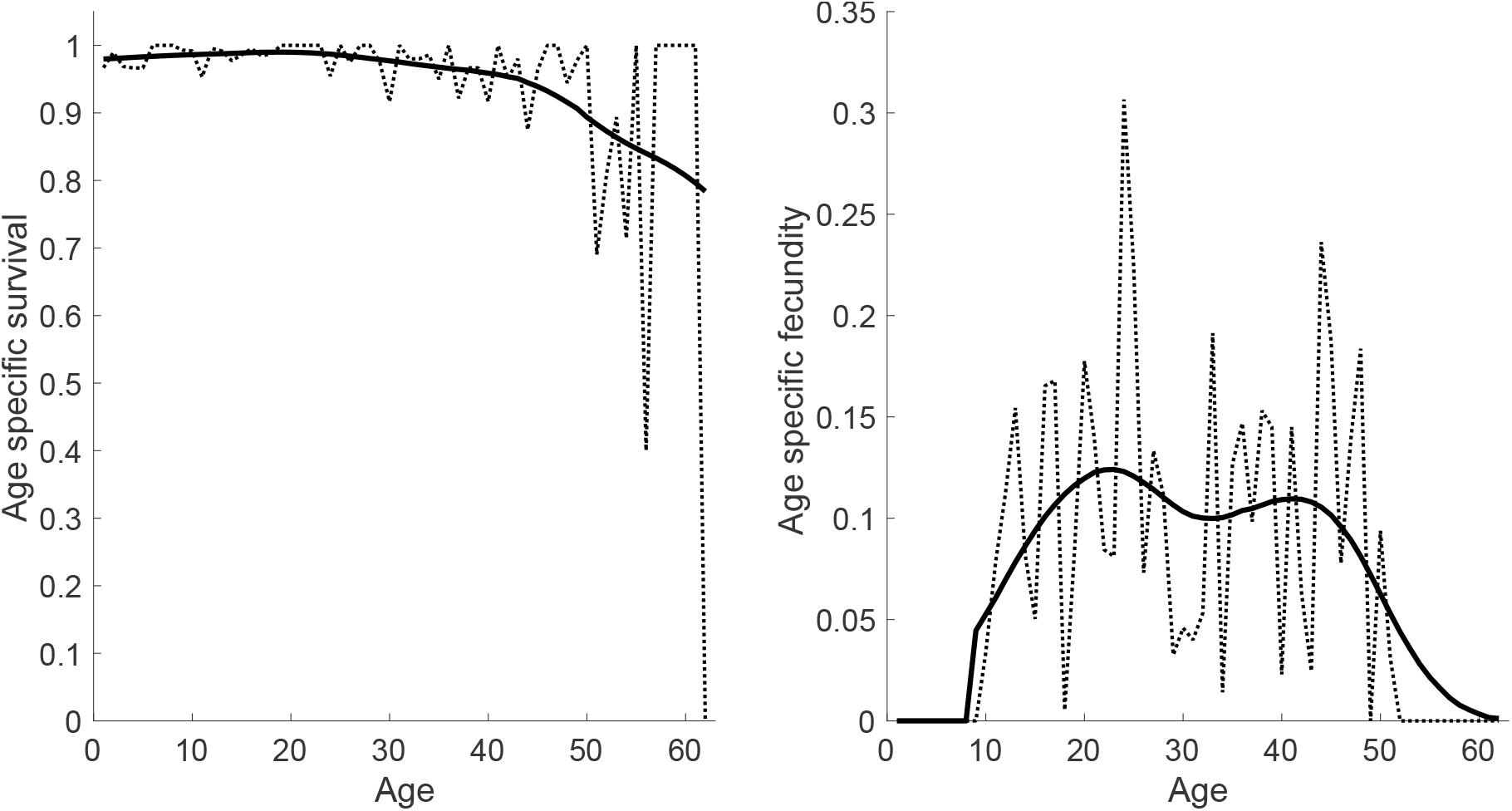
age-specific fertility and survival for African elephants used to parameterise our model. Dotted lines indicate the values reported by Wittemyer et al. (2021) and Wittemyer et al. (2013). Solid lines indicate the smoothed values used in this study calculated as the Gaussian-weighted moving average with a window of 25 year.

**Table S2:** Calculation of the proportional increase in fertility (***β***) due to the presence of a sister based on the fertility predicted by the statistical model from Lynch et al. (2019) with the statistical equation `logit(fertility)` ∼ `Sister` * `(Age-12)`, in which `Sister` is the absence (0) or presence (1) of at least one sister and `Age` is the age of the individual

| Age group | Predicted fertility<br>without sister | Predicted fertility<br>with sister | Proportional<br>increase used in<br>our model ($\beta$) |
| --- | --- | --- | --- |
| 9 | 0.0068 | 0.0202 | 1.9636 |
| 10 | 0.0086 | 0.0230 | 1.6788 |
| 11 | 0.0108 | 0.0261 | 1.4215 |
| 12 | 0.0135 | 0.0296 | 1.1893 |
| 13 | 0.0170 | 0.0336 | 0.9797 |
| 14 | 0.0212 | 0.0381 | 0.7908 |
| 15 | 0.0266 | 0.0431 | 0.6208 |
| 16 | 0.0331 | 0.0488 | 0.4678 |
| 17 | 0.0415 | 0.0552 | 0.3305 |
| 18 | 0.0517 | 0.0624 | 0.2076 |
| 19 | 0.0642 | 0.0707 | 0.0978 |
| 20 | 0.0794 | 0.0794 | 0.0000 |

**Figure S2:**
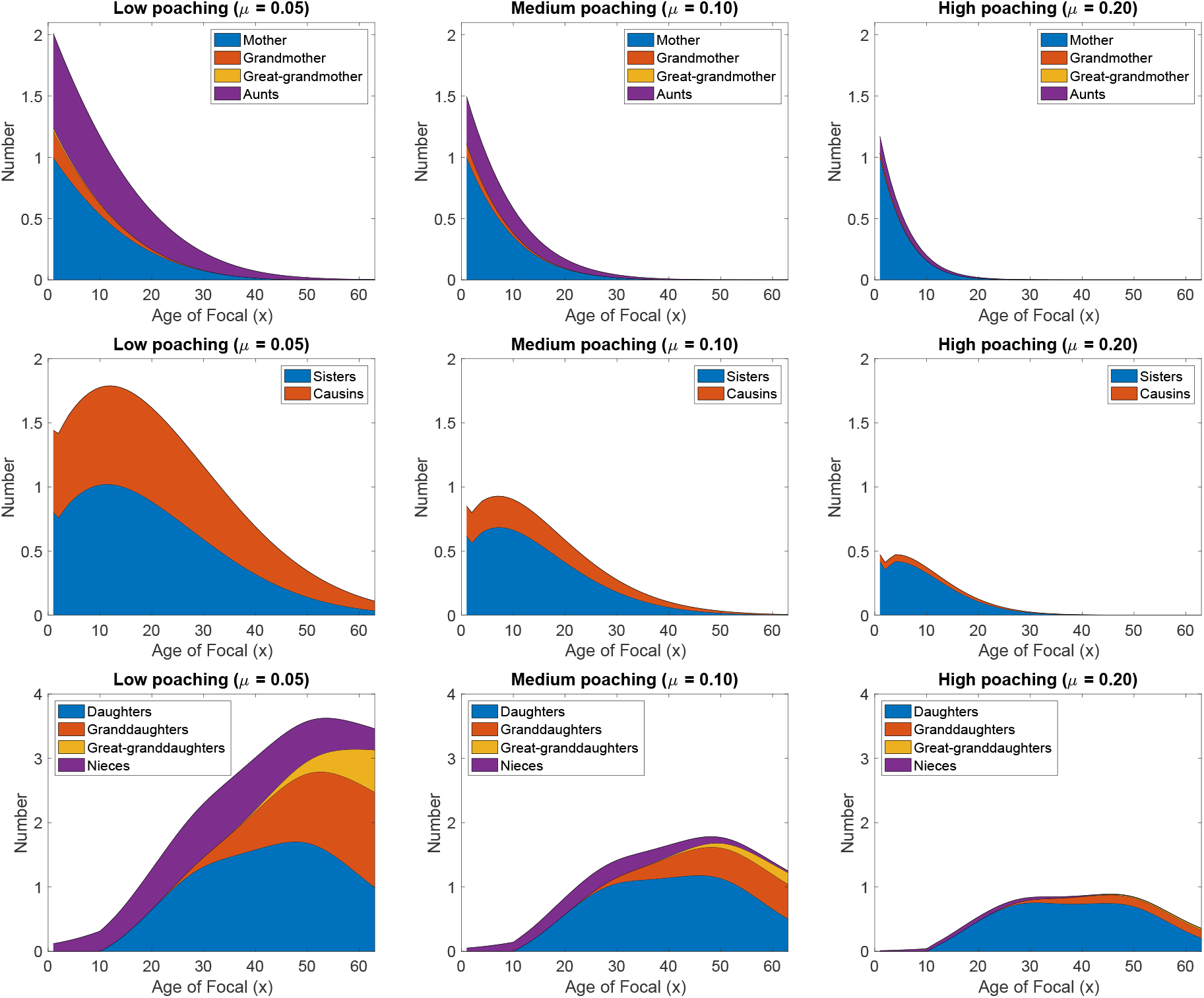
Expected number of kin throughout the life of a Focal individual for three different poaching intensities, while accounting for the interaction between the presence of the mother and juvenile survival.

**Figure S3:**
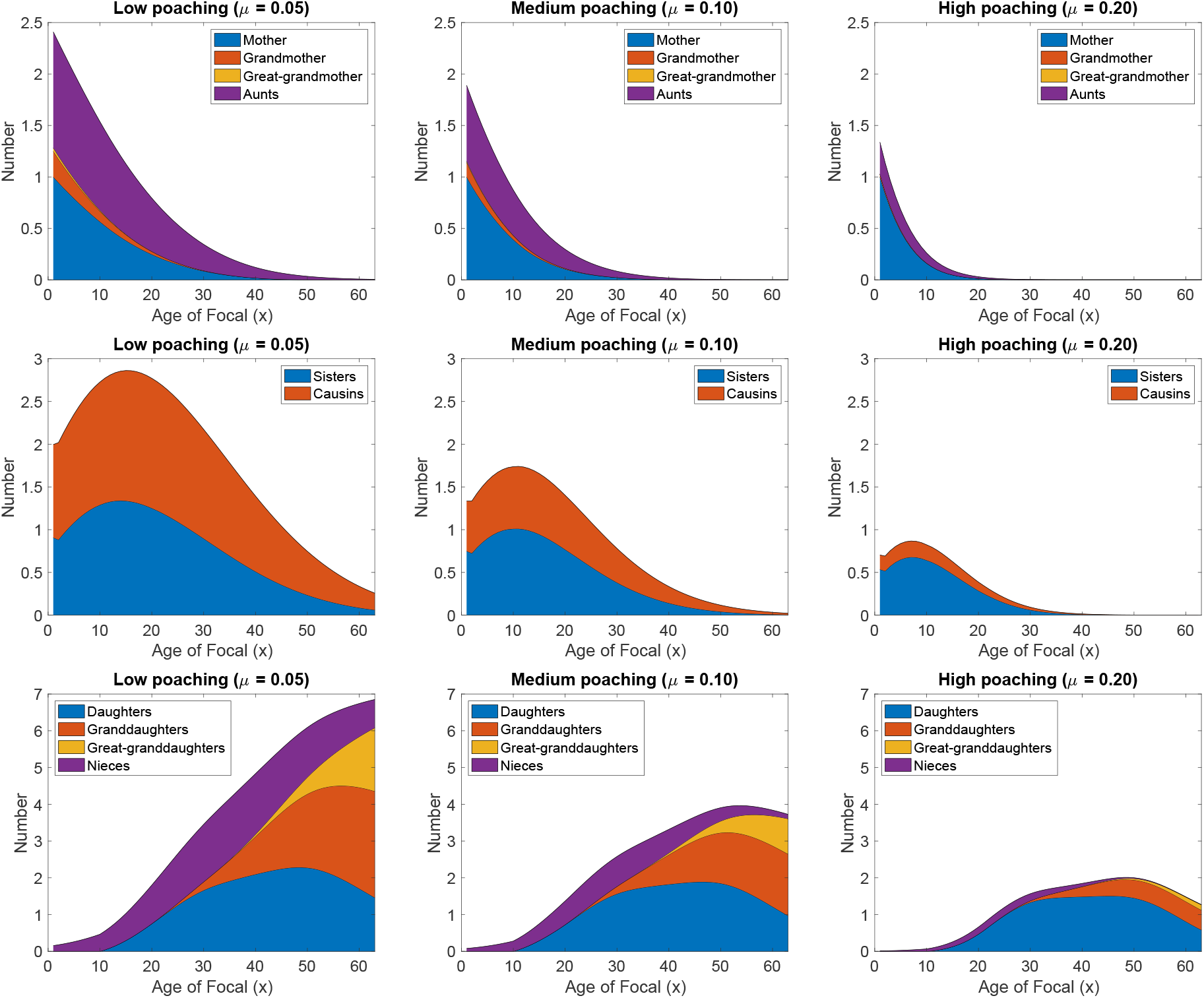
Expected number of kin throughout the life of a Focal individual for three different poaching intensities, while accounting for the interaction between the probability of having at least one sister and fertility of young females.

**Figure S4:**
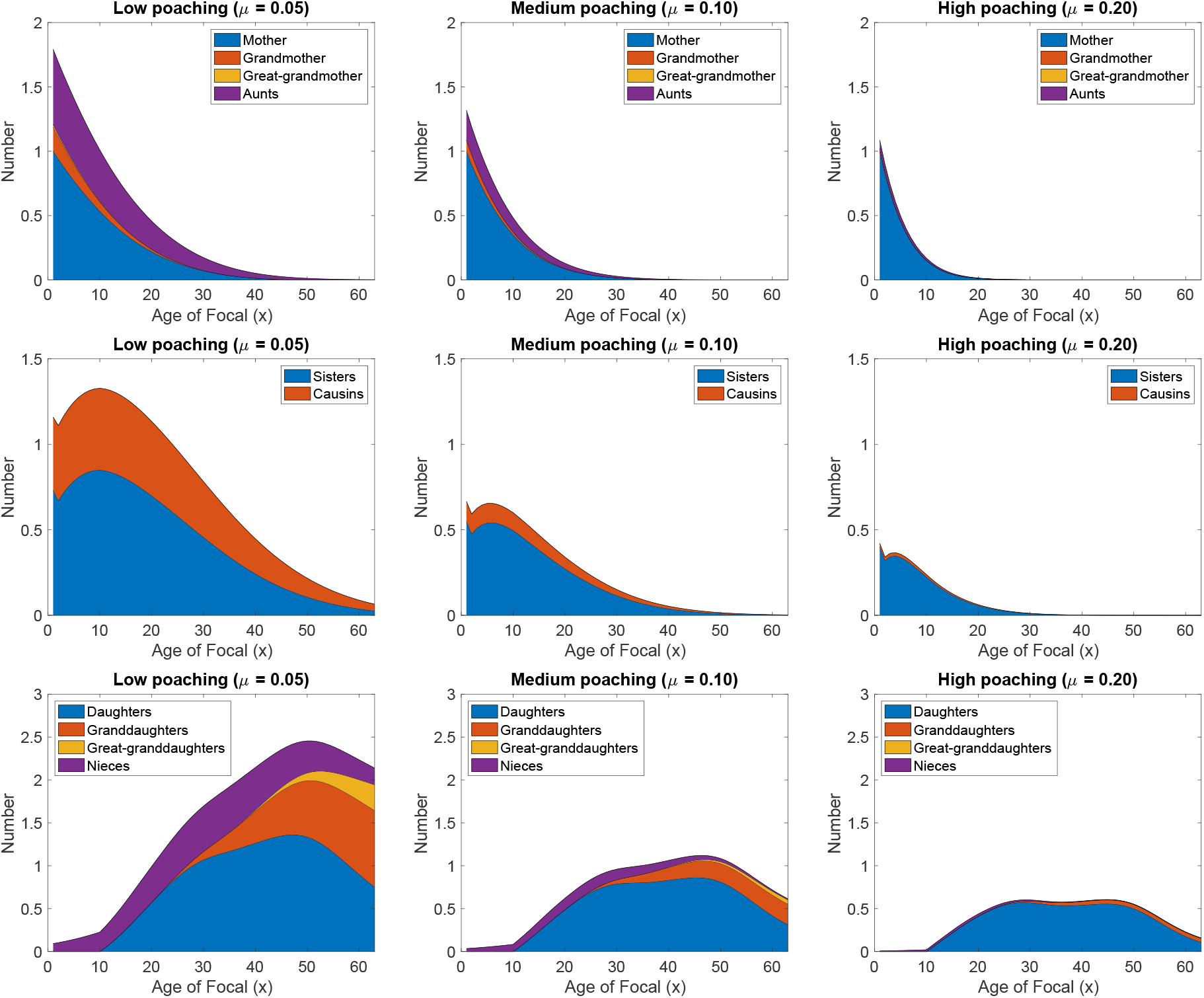
Expected number of kin throughout the life of a Focal individual for three different poaching intensities, while accounting for the interaction between the matriarch age and survival.

## Notes

### Competing Interest Statement

The authors have declared no competing interest.

### Summary of Updates

The model description has been rewritten to separate the general framework from the elephant specific example. The analysis for African elephants is re-run with a Bernoulli distribution for reproduction instead of a Poisson distribution. Several paragraphs are rewritten or added to the introduction and the discussion sections.

