## Supplementary material for "Family matters: Linking population growth, kin interactions, and African elephant social groups": Matlab code to run the model: c1_calcX.m

```
function X=c1_calcX(allkin)
% This function calculates the probability of a mother present for an an
% individual with ages 2 to 19 years for a given kin network.
%
% allkin is the full kin network as generated by the kinship function in
% the kinship_function.m file.

 

X = sum(allkin(:,2:19,5));

end
```
