## Supplementary material for "Family matters: Linking population growth, kin interactions, and African elephant social groups": Matlab code to run the model: c2_calcX.m

```
function X=c2_calcX(allkin)
% This function calculates the probability that an individual has at least
% one sisters at the ages 10 to 21 for a given kin network.
%
% allkin is the full kin network as generated by the kinship function in
% the kinship_function.m file.

upsilon = sum(allkin(:, 10:21,8))+sum(allkin(:, 10:21,9));
X = 1-exp(-upsilon);

end
```
