## Supplementary material for "Family matters: Linking population growth, kin interactions, and African elephant social groups": Matlab code to run the model: c3_calcX.m

```
function X=c3_calcX(allkin)
% This function calculates the expected oldest age in the family network at
% the ages 2 to 9 for a given kin network.
%
% allkin is the full kin network as generated by the kinship function in
% the kinship_function.m file.

% combine age structure of kin
z = sum(allkin(:,2:9,2:end),3);

% make empty array for results
Py = zeros(63,8);

% loop through ages
for x = 1:8

    % calculate probability when focal is the oldest individual
    Py(x,x) = ( z(x,x) / sum(z(x:end,x)))^sum(z(x:end,x));

    % calculate the probability that another age is the oldest age
    for i = (x+1):63
        Py(i,x) = (sum(z(x:i,x)) / sum(z(x:end,x)))^sum(z(x:end,x)) - (sum(z(x:i-1,x)) / sum(z(x:end,x)))^sum(z(x:end,x));
    end

end

% calculate the expected oldest age
X = (1:63)*Py;
```
