## Supplementary material for "Family matters: Linking population growth, kin interactions, and African elephant social groups": Matlab code to run the model: c4_calcX.m

```
function X=c4_calcX(allkin)
% this function combines all properties from all feedback from the research
% calculated with the c1_calcX, c2_calcX and c3_calcX
%
% allkin is the full kin network as generated by the kinship function in
% the kinship_function.m file.


% probability of having a mother at age 2 to 19
Xc1 = c1_calcX(allkin);

% probability of having at least one sister at age 10 to 21
Xc2 = c2_calcX(allkin);

% oldest age in the kin network at age 2 to 9
Xc3 = c3_calcX(allkin);

% combine values
X = [Xc1 Xc2 Xc3];

end
```
