## Supplementary material for "Family matters: Linking population growth, kin interactions, and African elephant social groups": Matlab code to run the model: kinship_function.m

```
function allkin=kinship_function(U,F)
% This function calculates the full family structure for all ages of focal.
% U = a square matrix with survival probabilities on the sub diagonal
% F = a square matrix with fecundity values in the first row
%
% the output of the function is a three dimentional array with the age structure of every
% family time of every age of focal.

% get number of age classes
[om,~]=size(U);

% make population projection matrix
Amat=U+F;

% get the normalized stable age distribution
[w,d]=eig(Amat);
d=diag(d);
pick=find(d==max(d));
w=w(:,pick)/sum(w(:,pick));

% create a transition matrix conditioned on survival
Usurv = U;
Usurv(Usurv ~= 0) = 1;

% create empty arrays for results
PHI = zeros(om,2*om); % Focal individual
A = zeros(om,2*om); % Daughters
B = zeros(om,2*om); % Granddaughters
C = zeros(om,2*om); % Greatgranddaughters
D = zeros(om,2*om); % Mothers
G = zeros(om,2*om); % Grandmothers
H = zeros(om,2*om); % Greatgrandmothers
M = zeros(om,2*om); % Older sisters
N = zeros(om,2*om); % Younger sisters
P = zeros(om,2*om); % Nieces through older sisters
Q = zeros(om,2*om); % Nieces through younger sisters
R = zeros(om,2*om); % Aunts older than mother
S = zeros(om,2*om); % Aunts youngher than mother
T = zeros(om,2*om); % Causins from older aunts
V = zeros(om,2*om); % Causins from youngher aunts

% the order of the following calculatings is chosen to minimize the number
% of for loops in the code. 

% We first project for focal, daughters, granddaughters,
% greatgranddaughters, mother, younger sisters and nieces through younger
% sisters

% intial conditions
D(:,1) = F(1,:)'.*w; 
D(:,1) = D(:,1)/sum(D(:,1));

% first timestep
PHI(1,2) = 1;
D(:,2) = U*D(:,1);

% Poisson N(:,2) = 0.5*(sum(F*D(:,1))/(1-exp(-sum(F*D(:,1))))-1);


% further projection
for ix=3:2*om
  PHI(:,ix) = Usurv*PHI(:,ix-1);
  A(:,ix) = U*A(:,ix-1) + F*PHI(:,ix-1);
  B(:,ix) = U*B(:,ix-1) + F*A(:,ix-1);
  C(:,ix) = U*C(:,ix-1) + F*B(:,ix-1);
  D(:,ix) = U*D(:,ix-1);
  N(:,ix) = U*N(:,ix-1) + F*D(:,ix-1);
  Q(:,ix) = U*Q(:,ix-1) + F*N(:,ix-1);
end

% we now project grandmothers, older sisters, nieces through older
% sisters, aunts younger than the mother and causins from younger aunts

% initial conditions
G(:,1) = D(:,2:om+1)*D(:,1);
M(:,1) = A(:,2:om+1)*D(:,1); 
P(:,1) = B(:,2:om+1)*D(:,1);
S(:,1) = N(:,2:om+1)*D(:,1);
V(:,1) = Q(:,2:om+1)*D(:,1);

% first timestep
G(:,2) = U*G(:,1);
M(:,2) = U*M(:,1); 

% Poisson M(:,2) = U*M(:,1) + 0.5*(sum(F*D(:,1))/(1-exp(-sum(F*D(:,1))))-1);

P(:,2) = U*P(:,1) + F*M(:,1);
S(:,2) = U*S(:,1) + F*G(:,1);
V(:,2) = U*V(:,1) + F*S(:,1);

% further projection
for ix=3:2*om
    G(:,ix) = U*G(:,ix-1);
    M(:,ix) = U*M(:,ix-1);
    P(:,ix) = U*P(:,ix-1) + F*M(:,ix-1);
    S(:,ix) = U*S(:,ix-1) + F*G(:,ix-1);
    V(:,ix) = U*V(:,ix-1) + F*S(:,ix-1);
end

% we now project grandmothers, aunts older than mother and causins from
% older aunts

% intial conditions
H(:,1) = G(:,2:om+1)*D(:,1);
R(:,1) = M(:,2:om+1)*D(:,1);
T(:,1) = P(:,2:om+1)*D(:,1);

% projection
for ix=2:2*om
    H(:,ix) = U*H(:,ix-1);
    R(:,ix) = U*R(:,ix-1);
    T(:,ix) = U*T(:,ix-1) + F*R(:,ix-1);
end

% combine all arrays with kin
allkin=cat(3,PHI,A,B,C,D,G,H,M,N,P,Q,R,S,T,V);

end
```
