## Supplementary material for "Family matters: Linking population growth, kin interactions, and African elephant social groups": Matlab code to run the model: Main_elephant_kin_interactions.m

```
%% PREABLE
% This matlab file contains the code to run the model and produce the
% figures for the "interactions in kinship networks" paper. 
%
% The following matlab files contain functions nessesairy to run the code
% in this file:
% - kinship_function.m; kinship_function(Umat, Fmat)
% - relatedness_high_function.m; relatedness_high_function(kinstruc)
% - relatedness_low_function.m; relatedness_low_function(kinstruc)
% - solveXkin.m; solveXkin(X,Fmat,Umatc1,@calcX)
% - c1_calcX.m; c1_calcX(kinstruc)
% - c2_calcX.m; c2_calcX(kinstruc)
% - c3_calcX.m; c3_calcX(kinstruc)
% - c4_calcX.m; c4_calcX(kinstruc)

%% GRAPHIC SETTINGS
% This sections sets the defaults graphic settings for the figures.

set(0,'defaultaxeslinewidth','factory');
set(0,'defaultaxesfontsize',18);
set(0,'DefaultAxesLinestyleOrder','-|--|:');
set(0,'DefaultLineLineWidth',3)

c0_0_style = "--black";
c0_colour = [0 0.4470 0.7410];
c1_colour = [0.8500 0.3250 0.0980];
c2_colour = [0.9290 0.6940 0.1250];
c3_colour = [0.4940 0.1840 0.5560];
c4_colour = "black";

%% LOAD LIFE HISTORY DATA
% This section loads the life history data used in the Afrikan elephant
% example.

% survival under low poaching from Wittemyer et. al. (2021)
p_wittemyer21 = [0.966886849,0.989374262,0.968764569,0.966482684,0.966562173,1,1,1,0.993006993,0.991735537,0.953030303,0.995215311,0.991735537,0.977272727,0.985930736,0.994949495,0.98474253,0.987012987,1,1,1,1,1,0.954545455,1,0.975,1,1,0.975,0.916666667,1,0.98,0.98,0.985714286,0.95,1,0.921666667,0.966666667,0.966666667,0.916666667,1,0.95,0.98,0.875,0.962962963,1,1,0.944444444,0.977777778,1]';

% fertility under low poaching from  Wittemyer et. al. (2021)
f_wittemyer21 = [0,0,0,0,0,0,0,0,0,0.033740865,0.078312597,0.112481964,0.154422837,0.081897711,0.050260092,0.165316894,0.16788314,0.005278144,0.084415584,0.177489177,0.139393939,0.084415584,0.081168831,0.30661157,0.223333333,0.073125,0.133333333,0.111666667,0.0325,0.045833333,0.04,0.0532,0.191333333,0.014081633,0.126666667,0.146969697,0.098311111,0.153055556,0.145,0.022916667,0.145,0.063333333,0.0245,0.23625,0.187242798,0.077777778,0.138888889,0.183641975,0,0.09375,0.030926145]';

% survivalship data for individuals age 51 to 63 from Wittemyer et. al.
% (2013)
l_wittemyer13 = [0.084 0.058 0.047 0.042 0.030 0.030 0.012 0.012 0.012 0.012 0.012 0.012 0.000]';


% convert survalship to survival
p_wittemyer13 =  l_wittemyer13(2:end)./l_wittemyer13(1:end-1);

% concatonate and smooth survival data
p_smooth = smoothdata([p_wittemyer21; p_wittemyer13],'gaussian',25);


% The reproductive success of indivuduals age 51 to 63 is expected to be
% zero
f_wittemyer13 = repelem(0,length(p_smooth) - length(f_wittemyer21))';

% concatonate and smooth fertility data
f_smooth = smoothdata([f_wittemyer21; f_wittemyer13],'gaussian',25);

% set fertility of juvenile classes back to zero
f_smooth(1:8) = 0;


%% PLOT LIFE HISTORY RATES
% this section plots the life history rates used for our simulations.
% Figure S1

lifehis_fig = figure('name','lifehis_fig');

tiledlayout(1,2)

% survival
nexttile
hold on
plot(p_smooth, '-black','LineWidth',3)
plot([p_wittemyer21; p_wittemyer13],':black','LineWidth',2)
hold off
xlim([0 63])
ylim([0.0 1.05])
xlabel('Age')
ylabel('Age specific survival')

% fecundity
nexttile
hold on
plot(f_smooth, '-black','LineWidth',3)
plot([f_wittemyer21; f_wittemyer13],':black','LineWidth',2)
hold off
xlim([0 63])
xlabel('Age')
ylabel('Age specific fecundity')

% save plot
lifehis_fig.Units = 'centimeters';
lifehis_fig.Position = [0 0 36 18];

exportgraphics(lifehis_fig,'figS1.pdf','Resolution',600)


%% INCLUDE ILLIGAL KILLING
% Here we include additional mortality to the survival values.

syms mup; % poaching mortality scalar
mupvalues = 0:0.002:0.2; % values for simulations

% age specific poaching sensitivity
rho = zeros(62,1);
rho(9:18) = 0.5;
rho(19:end) = 1;

% survival with poaching
p_poach = (1-rho*mup).*p_smooth;


%% MATRICES
% Here the default matrices without interactions are created.

Umat0 = diag(p_poach, -1 );

Fmat0 = zeros(63);
Fmat0(1, 1:end-1) = f_smooth;


%% POPULATION GROWTH WITHOUT FEEDBACK 
% Here we calculate the population growth without family feedback

% make empty arrays for results
c0_lambdas = zeros(1,101);
c0_eigvec = zeros(63,101);
c0_poprel_low = zeros(1,101);
c0_poprel_high = zeros(1,101);

% loop through the poaching pressures
for i = 1:length(mupvalues)

    % set poaching pressure in matrices
    mup = mupvalues(i);
    Amat = double(subs(Umat0)) + Fmat0;

    % calculate gpopulation rowthrate and structure
    [c0_eigvec(:,i), c0_lambdas(i)]  = eigs(Amat,1);
    c0_eigvec(:,i) = c0_eigvec(:,i)/sum(c0_eigvec(:,i));
    
    % calculate kin network 
    kinstruc_temp = kinship_function(double(subs(Umat0)),Fmat0);
    
    % calculate relatedness to kin network
    related_high_temp = relatedness_high_function(kinstruc_temp);
    related_low_temp = relatedness_low_function(kinstruc_temp);

    % weigh relatedness to population structure
    c0_poprel_low(i) = related_low_temp(1:63)*c0_eigvec(:,i);
    c0_poprel_high(i) = related_high_temp(1:63)*c0_eigvec(:,i);

    % report progress
    fprintf('mup=%f, progress %f\n', mup, i/101);
end

% set kin structure without poaching as baseline values

mup = 0;
kinstruc_base = kinship_function(double(subs(Umat0)),Fmat0);


%% PLOT RELATEDNESS AS FUNCTION OF AGE
% Here the relatedness as a function of age is plotted
% Figure 3

% Calculate the relatedness under four poaching pressures
mup = 0;
kinstruc_c0_00 = kinship_function(double(subs(Umat0)),Fmat0);
related_high_c0_00 = relatedness_high_function(kinstruc_c0_00);
related_low_c0_00 = relatedness_low_function(kinstruc_c0_00);

mup = 0.05;
kinstruc_c0_05 = kinship_function(double(subs(Umat0)),Fmat0);
related_high_c0_05 = relatedness_high_function(kinstruc_c0_05);
related_low_c0_05 = relatedness_low_function(kinstruc_c0_05);

mup = 0.1;
kinstruc_c0_10 = kinship_function(double(subs(Umat0)),Fmat0);
related_high_c0_10 = relatedness_high_function(kinstruc_c0_10);
related_low_c0_10 = relatedness_low_function(kinstruc_c0_10);

mup = 0.2;
kinstruc_c0_20 = kinship_function(double(subs(Umat0)),Fmat0);
related_high_c0_20 = relatedness_high_function(kinstruc_c0_20);
related_low_c0_20 = relatedness_low_function(kinstruc_c0_20);

% Create figure
c0_related_fig = figure('name','c0_related_fig');

tiledlayout(1,3)

% plot for mup=0
nexttile
hold on
patch([0:(length(related_low_c0_00)-1) fliplr(0:(length(related_low_c0_00)-1))], [related_high_c0_00  fliplr(related_low_c0_00)], c0_colour);
plot(0:(length(related_low_c0_00)-1), related_low_c0_00, 'Color', c0_colour)
plot(0:(length(related_high_c0_00)-1), related_high_c0_00, 'Color', c0_colour)
hold off 
alpha(0.5)
xlim([0 63])
ylim([0 0.5])
xlabel('Age of Focal (x)')
ylabel('Average ralatedness to kin (\eta)')
title("No poaching (\mu = 0.00)")

% plot for mup=0.1
nexttile
hold on
patch([0:(length(related_low_c0_10)-1) fliplr(0:(length(related_low_c0_10)-1))], [related_high_c0_10  fliplr(related_low_c0_10)], c0_colour);
plot(0:(length(related_low_c0_10)-1), related_low_c0_10, 'Color', c0_colour)
plot(0:(length(related_high_c0_10)-1), related_high_c0_10, 'Color', c0_colour)
hold off 
alpha(0.5)
xlim([0 63])
ylim([0 0.5])
xlabel('Age of Focal (x)')
ylabel('Average ralatedness to kin (\eta)')
title("Medium poaching (\mu = 0.10)")

% plot for mup=0.2
nexttile
hold on
patch([0:(length(related_low_c0_20)-1) fliplr(0:(length(related_low_c0_20)-1))], [related_high_c0_20  fliplr(related_low_c0_20)], c0_colour);
plot(0:(length(related_low_c0_20)-1), related_low_c0_20, 'Color', c0_colour)
plot(0:(length(related_high_c0_20)-1), related_high_c0_20, 'Color', c0_colour)
hold off 
alpha(0.5)
xlim([0 63])
ylim([0 0.5])
xlabel('Age of Focal (x)')
ylabel('Average ralatedness to kin (\eta)')
title("High poaching (\mu = 0.20)")

% save plot
c0_related_fig.Units = 'centimeters';
c0_related_fig.Position = [0 0 54 18];

exportgraphics(c0_related_fig,'fig3.pdf','Resolution',600)


%% EFFECT MOTHER ON JUVENILE SURVIVAL (c1)
% This section solves the kin network with the effect of the mother
% presence on juvenile survival

% reset values
syms mup;
X = sym('x',[1 18]);
Fmatc1 = Fmat0;
Umatc1 = Umat0;

% claculate baseline
c1Xbase = c1_calcX(kinstruc_base);

% Load values of interactions (Table S2)
alphax = zeros(63, 1);
alphax(1:2) = 1;
alphax(3:8) = 0.1427;
alphax(9:18) = 0.0387;

% Include interactions in the matrices
Umatc1(:,1:18) = (1-(c1Xbase - X).*alphax(1:18)').*Umat0(:,1:18);

% Make empty arrays for results
c1_lambdas = zeros(1,101);
c1_eigvec = zeros(63,101);
c1_Xsolve = zeros(1,18,101);
c1_poprel_low = zeros(1,101);
c1_poprel_high = zeros(1,101);

% set values without poaching
c1_lambdas(1) = c0_lambdas(1);
c1_Xsolve(:,:,1) = c1Xbase;

c1_poprel_low(1) = c0_poprel_low(1);
c1_poprel_high(1) = c0_poprel_high(1);

% loop through the poaching pressures
for i = 2:length(mupvalues)

    % set poaching pressure
    mup = mupvalues(i);

    % solve for kin network
    c1_Xsolve(:,:,i) = fsolve(@(X)solveXkin(X,Fmatc1,subs(Umatc1),@c1_calcX),c1_Xsolve(:,:,i-1),optimoptions('fsolve','Display','iter','UseParallel',false));
    
    % calculate population porjection matrix
    Amatc1 = double(subs(subs(Umatc1,X,c1_Xsolve(:,:,i)))) + Fmatc1;

    % calculate population growthrate and structure
    [c1_eigvec(:,i), c1_lambdas(i)]  = eigs(Amatc1,1);
    c1_eigvec(:,i) = c1_eigvec(:,i)/sum(c1_eigvec(:,i));

    % calculate kinstructure
    kinstruc_temp = kinship_function( double(subs(subs(Umatc1,X,c1_Xsolve(:,:,i)))),Fmatc1);
    
    % calculate relatedness
    related_high_temp = relatedness_high_function(kinstruc_temp);
    related_low_temp = relatedness_low_function(kinstruc_temp);

    c1_poprel_low(i) = related_low_temp(1:63)*c1_eigvec(:,i);
    c1_poprel_high(i) = related_high_temp(1:63)*c1_eigvec(:,i);

    % report progress
    fprintf('\n\n\nSucceeded, mup=%f, lambda = %f, progress %f\n', mup, c1_lambdas(i), i/101);
end


%% PLOT MOTHER EFFECT ON JUVENILE SURVIVAL FIGURE
% here we plot the probability of a mother present under the effect of
% mothers on the juvenile survival for several poaching pressures.
% Figure 5

% Calculate kin structure
mup = 0;
kinstruc_c1_00 = kinship_function(double(subs(subs(Umatc1,X,c1_Xsolve(:,:,1)))),Fmatc1);
related_high_c1_00 = relatedness_high_function(kinstruc_c1_00);
related_low_c1_00 = relatedness_low_function(kinstruc_c1_00);

mup = 0.05;
kinstruc_c1_05 = kinship_function(double(subs(subs(Umatc1,X,c1_Xsolve(:,:,26)))),Fmatc1);
related_high_c1_05 = relatedness_high_function(kinstruc_c1_05);
related_low_c1_05 = relatedness_low_function(kinstruc_c1_05);

mup = 0.1;
kinstruc_c1_10 = kinship_function(double(subs(subs(Umatc1,X,c1_Xsolve(:,:,51)))),Fmatc1);
related_high_c1_10 = relatedness_high_function(kinstruc_c1_10);
related_low_c1_10 = relatedness_low_function(kinstruc_c1_10);

mup = 0.2;
kinstruc_c1_20 = kinship_function(double(subs(subs(Umatc1,X,c1_Xsolve(:,:,101)))),Fmatc1);
related_high_c1_20 = relatedness_high_function(kinstruc_c1_20);
related_low_c1_20 = relatedness_low_function(kinstruc_c1_20);

% calculate probability of presence of a mother
mother_c0_00 = sum(kinstruc_c0_00(:,:,5));
mother_c0_05 = sum(kinstruc_c0_05(:,:,5));
mother_c0_10 = sum(kinstruc_c0_10(:,:,5));
mother_c0_20 = sum(kinstruc_c0_20(:,:,5));

mother_c1_00 = sum(kinstruc_c1_00(:,:,5));
mother_c1_05 = sum(kinstruc_c1_05(:,:,5));
mother_c1_10 = sum(kinstruc_c1_10(:,:,5));
mother_c1_20 = sum(kinstruc_c1_20(:,:,5));

% create figure
c1_mother_fig = figure('name','c1_mother_fig');

tiledlayout(1,3)

% mup = 0.05
nexttile
hold on
plot(0:(length(mother_c0_00)-1), mother_c0_00, c0_0_style )
plot(0:(length(mother_c0_05)-1), mother_c0_05, 'Color', c0_colour)
plot(0:(length(mother_c1_05)-1), mother_c1_05, 'Color', c1_colour)
hold off 
xlim([0 63])
legend('No poaching','No feedback','Effect of mothers on survival')
xlabel('Age of Focal (x)')
ylabel('Mother')
title("Low poaching (\mu = 0.05)")

% mup = 0.1
nexttile
hold on
plot(0:(length(mother_c0_00)-1), mother_c0_00, c0_0_style )
plot(0:(length(mother_c0_10)-1), mother_c0_10, 'Color', c0_colour)
plot(0:(length(mother_c1_10)-1), mother_c1_10, 'Color', c1_colour)
hold off 
xlim([0 63])
legend('No poaching','No feedback','Effect of mothers on survival')
xlabel('Age of Focal (x)')
ylabel('Mother')
title("Medium poaching (\mu = 0.10)")

% mup = 0.2
nexttile
hold on
plot(0:(length(mother_c0_00)-1), mother_c0_00, c0_0_style )
plot(0:(length(mother_c0_20)-1), mother_c0_20, 'Color', c0_colour)
plot(0:(length(mother_c1_20)-1), mother_c1_20, 'Color', c1_colour)
hold off 
xlim([0 63])
legend('No poaching','No feedback','Effect of mothers on survival')
xlabel('Age of Focal (x)')
ylabel('Mother')
title("High poaching (\mu = 0.20)")

% set figure size
c1_mother_fig.Units = 'centimeters';
c1_mother_fig.Position = [0 0 54 18];

% save figure 
exportgraphics(c1_mother_fig,'fig5.pdf','Resolution',600)


%% PLOT MOTHER EFFECT  ON SURVIVAL FULL KINSTRUCTURE
% Here the full kin sturcture with the effect of mothers on juvenile
% survival is plotted
% Figure S2

% rotate kinstructure
total_kinstruc_c1_05 = shiftdim(sum(kinstruc_c1_05),1);
total_kinstruc_c1_10 = shiftdim(sum(kinstruc_c1_10),1);
total_kinstruc_c1_20 = shiftdim(sum(kinstruc_c1_20),1);

% create figure
c1_totkin_fig = figure('name','c1_totkin_fig');

tiledlayout(3,3)

% mup = 0.05 old generation
nexttile
area([total_kinstruc_c1_05(:,[5 6 7]) sum(total_kinstruc_c1_05(:,[12 13]),2) ] ) 
xlim([0 63])
ylim([0 2.1])
legend('Mother','Grandmother','Great-grandmother','Aunts')
xlabel('Age of Focal (x)')
ylabel('Number')
title("Low poaching (\mu = 0.05)")

% mup = 0.1 old generation
nexttile
area([total_kinstruc_c1_10(:,[5 6 7]) sum(total_kinstruc_c1_10(:,[12 13]),2) ] ) 
xlim([0 63])
ylim([0 2.1])
legend('Mother','Grandmother','Great-grandmother','Aunts')
xlabel('Age of Focal (x)')
ylabel('Number')
title("Medium poaching (\mu = 0.10)")

% mup = 0.2 old generation
nexttile
area([total_kinstruc_c1_20(:,[5 6 7]) sum(total_kinstruc_c1_20(:,[12 13]),2) ] ) 
xlim([0 63])
ylim([0 2.1])
legend('Mother','Grandmother','Great-grandmother','Aunts')
xlabel('Age of Focal (x)')
ylabel('Number')
title("High poaching (\mu = 0.20)")

% mup = 0.05 middle generation
nexttile
area([sum(total_kinstruc_c1_05(:,[8 9]),2) sum(total_kinstruc_c0_05(:,[14 15]),2) ] ) 
xlim([0 63])
ylim([0 2])
legend('Sisters','Causins')
xlabel('Age of Focal (x)')
ylabel('Number')
title("Low poaching (\mu = 0.05)")

% mup = 0.1 middle generation
nexttile
area([sum(total_kinstruc_c0_10(:,[8 9]),2) sum(total_kinstruc_c1_10(:,[14 15]),2) ] ) 
xlim([0 63])
ylim([0 2])
legend('Sisters','Causins')
xlabel('Age of Focal (x)')
ylabel('Number')
title("Medium poaching (\mu = 0.10)")

% mup = 0.2 middle generation
nexttile
area([sum(total_kinstruc_c1_20(:,[8 9]),2) sum(total_kinstruc_c1_20(:,[14 15]),2) ] ) 
xlim([0 63])
ylim([0 2])
legend('Sisters','Causins')
xlabel('Age of Focal (x)')
ylabel('Number')
title("High poaching (\mu = 0.20)")

% mup = 0.05 young generation
nexttile
area([total_kinstruc_c1_05(:,[2 3 4]) sum(total_kinstruc_c1_05(:,[10 11]),2) ] ) 
xlim([0 63])
ylim([0 4])
legend('Daughters','Granddaughters', 'Great-granddaughters',"Nieces",'Location','northwest')
xlabel('Age of Focal (x)')
ylabel('Number')
title("Low poaching (\mu = 0.05)")

% mup = 0.1 young generation
nexttile
area([total_kinstruc_c1_10(:,[2 3 4]) sum(total_kinstruc_c1_10(:,[10 11]),2) ] ) 
xlim([0 63])
ylim([0 4])
legend('Daughters','Granddaughters', 'Great-granddaughters',"Nieces",'Location','northwest')
xlabel('Age of Focal (x)')
ylabel('Number')
title("Medium poaching (\mu = 0.10)")

% mup = 0.2 young generation
nexttile
area([total_kinstruc_c1_20(:,[2 3 4]) sum(total_kinstruc_c1_20(:,[10 11]),2) ] ) 
xlim([0 63])
ylim([0 4])
legend('Daughters','Granddaughters', 'Great-granddaughters',"Nieces",'Location','northwest')
xlabel('Age of Focal (x)')
ylabel('Number')
title("High poaching (\mu = 0.20)")

% set figure size
c1_totkin_fig.Units = 'centimeters';
c1_totkin_fig.Position = [0 0 54 70];

% save figure
exportgraphics(c1_totkin_fig,'figS2','Resolution',600)


%% EFFECT SISTERS ON FECUNDITY (c2)
% This section solves the kin network with the effect of the presence of a
% sister on the fecundity of females.

% reset values
syms mup
Fmatc2 = sym(Fmat0);
Umatc2 = Umat0;
X = sym('x',[1, length(c2Xbase)]);

% calculate baseline
c2Xbase = c2_calcX(kinstruc_base);

% Load values of interactions (Table S2)
Lynch_pred_nosister = exp(-4.29 + 0.23*((9:20)-12)) ./ (1 + exp(-4.29 + 0.23*((9:20)-12)));
Lynch_pred_sister = exp(-4.29 + 0.23*((9:20)-12) + 0.8 - 0.1*((9:20)-12)) ./ (1 + exp(-4.29 + 0.23*((9:20)-12) + 0.8 - 0.1*((9:20)-12)));

kappa = (Lynch_pred_sister - Lynch_pred_nosister)./Lynch_pred_nosister;

% Include interaction in the matrices
Fmatc2(:,9:20) = (1 + ( X - c2Xbase ) .* kappa) .* Fmat0(:,9:20);


% create empty arrays
c2_lambdas = zeros(1,101);
c2_eigvec = zeros(63,101);
c2_Xsolve = zeros(1,length(c2Xbase),101);
c2_poprel_low = zeros(1,101);
c2_poprel_high = zeros(1,101);

% set values without poaching
c2_lambdas(1) = c0_lambdas(1);
c2_Xsolve(:,:,1) = c2Xbase;

c2_poprel_low(1) = c0_poprel_low(1);
c2_poprel_high(1) = c0_poprel_high(1);


% loop through poaching pressures
for i = 2:length(mupvalues)
    
    % set poaching pressure
    mup = mupvalues(i);

    % solve kin stucture
    c2_Xsolve(:,:,i) = fsolve(@(X)solveXkin(X,Fmatc2,subs(Umatc2),@c2_calcX),c2_Xsolve(:,:,i-1),optimoptions('fsolve','Display','iter','UseParallel',false));
    
    % create population projection matrix
    Amatc2 = double(subs(subs(Umatc2,X,c2_Xsolve(:,:,i))) + subs(subs(Fmatc2,X,c2_Xsolve(:,:,i))));
    
    % calcualte population growth rate and structure
    [c2_eigvec(:,i), c2_lambdas(i)]  = eigs(Amatc2,1);
    c2_eigvec(:,i) = c2_eigvec(:,i)/sum(c2_eigvec(:,i));

    % calculate kin network
    kinstruc_temp = kinship_function( double(subs(subs(Umatc2,X,c2_Xsolve(:,:,i)))),double(subs(subs(Fmatc2,X,c2_Xsolve(:,:,i)))));
    
    % calculate realtedness
    related_high_temp = relatedness_high_function(kinstruc_temp);
    related_low_temp = relatedness_low_function(kinstruc_temp);
    
    c2_poprel_low(i) = related_low_temp(1:63)*c2_eigvec(:,i);
    c2_poprel_high(i) = related_high_temp(1:63)*c2_eigvec(:,i);

    % report progress
    fprintf('\n\n\nSucceeded, mup=%f, lambda = %f, progress %f\n', mup, c2_lambdas(i), i/101);

end


%% PLOT SISTER EFFECT ON FECUNDITY FIGURE
% here we plot the probability of having at least one sister with the
% effect of sisters on fecundity
% Figure 6

% Calculate kin structure
mup = 0;
kinstruc_c2_00 = kinship_function(double(subs(Umatc2)),double(subs(subs(Fmatc2,X,c2_Xsolve(:,:,1)))));
related_high_c2_00 = relatedness_high_function(kinstruc_c2_00);
related_low_c2_00 = relatedness_low_function(kinstruc_c2_00);

mup = 0.05;
kinstruc_c2_05 = kinship_function(double(subs(Umatc2)),double(subs(subs(Fmatc2,X,c2_Xsolve(:,:,26)))));
related_high_c2_05 = relatedness_high_function(kinstruc_c2_05);
related_low_c2_05 = relatedness_low_function(kinstruc_c2_05);

mup = 0.10;
kinstruc_c2_10 = kinship_function(double(subs(Umatc2)),double(subs(subs(Fmatc2,X,c2_Xsolve(:,:,51)))));
related_high_c2_10 = relatedness_high_function(kinstruc_c2_10);
related_low_c2_10 = relatedness_low_function(kinstruc_c2_10);

mup = 0.20;
kinstruc_c2_20 = kinship_function(double(subs(Umatc2)),double(subs(subs(Fmatc2,X,c2_Xsolve(:,:,101)))));
related_high_c2_20 = relatedness_high_function(kinstruc_c2_20);
related_low_c2_20 = relatedness_low_function(kinstruc_c2_20);

% Calculate probability of at least one sister
sisters_c0_00 = 1-exp(-sum(kinstruc_c0_00(:,:,8)) - sum(kinstruc_c0_00(:,:,9)));
sisters_c0_05 = 1-exp(-sum(kinstruc_c0_05(:,:,8)) - sum(kinstruc_c0_05(:,:,9)));
sisters_c0_10 = 1-exp(-sum(kinstruc_c0_10(:,:,8)) - sum(kinstruc_c0_10(:,:,9)));
sisters_c0_20 = 1-exp(-sum(kinstruc_c0_20(:,:,8)) - sum(kinstruc_c0_20(:,:,9)));

sisters_c2_00 = 1-exp(-sum(kinstruc_c2_00(:,:,8)) - sum(kinstruc_c2_00(:,:,9)));
sisters_c2_05 = 1-exp(-sum(kinstruc_c2_05(:,:,8)) - sum(kinstruc_c2_05(:,:,9)));
sisters_c2_10 = 1-exp(-sum(kinstruc_c2_10(:,:,8)) - sum(kinstruc_c2_10(:,:,9)));
sisters_c2_20 = 1-exp(-sum(kinstruc_c2_20(:,:,8)) - sum(kinstruc_c2_20(:,:,9)));

% create figure
c2_sisters_fig = figure('name','c2_sisters_fig');

tiledlayout(1,3)

% mup = 0.05
nexttile
hold on
plot(0:(length(sisters_c0_00)-1), sisters_c0_00, c0_0_style )
plot(0:(length(sisters_c0_05)-1), sisters_c0_05, 'Color', c0_colour)
plot(0:(length(sisters_c2_05)-1), sisters_c2_05, 'Color', c2_colour)
yline(1)
hold off 
xlim([0 63])
legend('No poaching','No feedback','Effect of sisters on fecundity','')
xlabel('Age of Focal (x)')
ylabel('Probability of at least one sister')
title("Low poaching (\mu = 0.05)")

% mup = 0.1
nexttile
hold on
plot(0:(length(sisters_c0_00)-1), sisters_c0_00, c0_0_style )
plot(0:(length(sisters_c0_10)-1), sisters_c0_10, 'Color', c0_colour)
plot(0:(length(sisters_c2_10)-1), sisters_c2_10, 'Color', c2_colour)
yline(1)
hold off 
xlim([0 63])
legend('No poaching','No feedback','Effect of sisters on fecundity','')
xlabel('Age of Focal (x)')
ylabel('Probability of at least one sister')
title("Medium poaching (\mu = 0.10)")

% mup = 0.2
nexttile
hold on
plot(0:(length(sisters_c0_00)-1), sisters_c0_00, c0_0_style )
plot(0:(length(sisters_c0_20)-1), sisters_c0_20, 'Color', c0_colour)
plot(0:(length(sisters_c2_20)-1), sisters_c2_20, 'Color', c2_colour)
yline(1)
hold off 
xlim([0 63])
legend('No poaching','No feedback','Effect of sisters on fecundity','')
xlabel('Age of Focal (x)')
ylabel('Probability of at least one sister')
title("High poaching (\mu = 0.20)")

% set figure size
c2_sisters_fig.Units = 'centimeters';
c2_sisters_fig.Position = [0 0 54 18];

% save figure
exportgraphics(c2_sisters_fig,'fig6.pdf','Resolution',600)


%% PLOT SISTER EFFECT ON FECUNDITY FULL KINSTRUCTURE
% Here the full kin sturcture with the effect of sisters on the fecundity
% Figure S3

% rotate kinstructure
total_kinstruc_c2_05 = shiftdim(sum(kinstruc_c2_05),1);
total_kinstruc_c2_10 = shiftdim(sum(kinstruc_c2_10),1);
total_kinstruc_c2_20 = shiftdim(sum(kinstruc_c2_20),1);

% create figure
c2_totkin_fig = figure('name','c2_totkin_fig');

tiledlayout(3,3)

% mup=0.05; old generation
nexttile
area([total_kinstruc_c2_05(:,[5 6 7]) sum(total_kinstruc_c2_05(:,[12 13]),2) ] ) 
xlim([0 63])
ylim([0 2.5])
legend('Mother','Grandmother','Great-grandmother','Aunts')
xlabel('Age of Focal (x)')
ylabel('Number')
title("Low poaching (\mu = 0.05)")

% mup=0.1; old generation
nexttile
area([total_kinstruc_c2_10(:,[5 6 7]) sum(total_kinstruc_c2_10(:,[12 13]),2) ] ) 
xlim([0 63])
ylim([0 2.5])
legend('Mother','Grandmother','Great-grandmother','Aunts')
xlabel('Age of Focal (x)')
ylabel('Number')
title("Medium poaching (\mu = 0.10)")

% mup=0.2; old generation
nexttile
area([total_kinstruc_c2_20(:,[5 6 7]) sum(total_kinstruc_c2_20(:,[12 13]),2) ] ) 
xlim([0 63])
ylim([0 2.5])
legend('Mother','Grandmother','Great-grandmother','Aunts')
xlabel('Age of Focal (x)')
ylabel('Number')
title("High poaching (\mu = 0.20)")

% mup=0.05; middle generation
nexttile
area([sum(total_kinstruc_c2_05(:,[8 9]),2) sum(total_kinstruc_c2_05(:,[14 15]),2) ] ) 
xlim([0 63])
ylim([0 3])
legend('Sisters','Causins')
xlabel('Age of Focal (x)')
ylabel('Number')
title("Low poaching (\mu = 0.05)")

% mup=0.1; middle generation
nexttile
area([sum(total_kinstruc_c2_10(:,[8 9]),2) sum(total_kinstruc_c2_10(:,[14 15]),2) ] ) 
xlim([0 63])
ylim([0 3])
legend('Sisters','Causins')
xlabel('Age of Focal (x)')
ylabel('Number')
title("Medium poaching (\mu = 0.10)")

% mup=0.2; middle generation
nexttile
area([sum(total_kinstruc_c2_20(:,[8 9]),2) sum(total_kinstruc_c2_20(:,[14 15]),2) ] ) 
xlim([0 63])
ylim([0 3])
legend('Sisters','Causins')
xlabel('Age of Focal (x)')
ylabel('Number')
title("High poaching (\mu = 0.20)")

% mup=0.05; young generation
nexttile
area([total_kinstruc_c2_05(:,[2 3 4]) sum(total_kinstruc_c2_05(:,[10 11]),2) ] ) 
xlim([0 63])
ylim([0 7])
legend('Daughters','Granddaughters', 'Great-granddaughters',"Nieces",'Location','northwest')
xlabel('Age of Focal (x)')
ylabel('Number')
title("Low poaching (\mu = 0.05)")

% mup=0.1; young generation
nexttile
area([total_kinstruc_c2_10(:,[2 3 4]) sum(total_kinstruc_c2_10(:,[10 11]),2) ] ) 
xlim([0 63])
ylim([0 7])
legend('Daughters','Granddaughters', 'Great-granddaughters',"Nieces",'Location','northwest')
xlabel('Age of Focal (x)')
ylabel('Number')
title("Medium poaching (\mu = 0.10)")

% mup=0.2; young generation
nexttile
area([total_kinstruc_c2_20(:,[2 3 4]) sum(total_kinstruc_c2_20(:,[10 11]),2) ] ) 
xlim([0 63])
ylim([0 7])
legend('Daughters','Granddaughters', 'Great-granddaughters',"Nieces",'Location','northwest')
xlabel('Age of Focal (x)')
ylabel('Number')
title("High poaching (\mu = 0.20)")

% set figure size
c2_totkin_fig.Units = 'centimeters';
c2_totkin_fig.Position = [0 0 54 70];

% save figure
exportgraphics(c2_totkin_fig,'figS3.pdf','Resolution',600)


%% EFFECT MATRIARCH AGE ON SURVIVAL (c3)
% This section solves the kin network with the effect of oldest age on
% survival.

%reset values
syms mup
Fmatc3 = Fmat0;
Umatc3 = Umat0;
X = sym('x',[1, length(c3Xbase)]);

% calculate expected oldest age without poaching (X base)
c3Xbase = c3_calcX(kinstruc_base);

% Add effect to matrix
Umatc3(:,1:8) = Umat0(:,1:8) .* (1+exp(0.047.*c3Xbase)) ./ (1+exp(0.047.*X)) .* exp(0.047*(X-c3Xbase));

% make empty arrays for results
c3_lambdas = zeros(1,101);
c3_eigvec = zeros(63,101);

c3_Xsolve = zeros(1,length(c3Xbase),101);

c3_poprel_low = zeros(1,101);
c3_poprel_high = zeros(1,101);

% set values without poaching
c3_lambdas(1) = c0_lambdas(1);

c3_Xsolve(:,:,1) = c3Xbase;

c3_eigvec(:,1) = c0_eigvec(:,1);
c3_poprel_low(1) = c0_poprel_low(1);
c3_poprel_high(1) = c0_poprel_high(1);

% loop through poaching pressures
for i = 2:length(mupvalues)

    % set poaching pressure
    mup = mupvalues(i);
    
    % solve kin structure
    c3_Xsolve(:,:,i) = fsolve(@(X)solveXkin(X,Fmatc3,subs(Umatc3),@c3_calcX),c3_Xsolve(:,:,i-1),optimoptions('fsolve','Display','iter','UseParallel',false));

    % make population porjection matrix
    Amatc3 = double(subs(subs(Umatc3,X,c3_Xsolve(:,:,i))) + subs(subs(Fmatc3,X,c3_Xsolve(:,:,i))));
    
    % calculate population growth rate and structure
   [c3_eigvec(:,i), c3_lambdas(i)]  = eigs(Amatc3,1);
    c3_eigvec(:,i) = c3_eigvec(:,i)/sum(c3_eigvec(:,i));

    % calculate kinstructure
    kinstruc_temp = kinship_function( double(subs(subs(Umatc3,X,c3_Xsolve(:,:,i)))),double(subs(subs(Fmatc3,X,c3_Xsolve(:,:,i)))));
    
    % calculate relatedness
    related_high_temp = relatedness_high_function(kinstruc_temp);
    related_low_temp = relatedness_low_function(kinstruc_temp);

    c3_poprel_low(i) = related_low_temp(1:63)*c3_eigvec(:,i);
    c3_poprel_high(i) = related_high_temp(1:63)*c3_eigvec(:,i);

    % report progress
    fprintf('\n\n\nSucceeded, mup=%f, lambda = %f, progress %f\n', mup, c3_lambdas(i), i/101);
end


%% PLOT OLDEST AGE ON SURVIVAL FIGURE
% here we plot the expected matriarch age when including the effect of
% matriarch age in survival
% Figure 7

% Calculate kinstructure for four different poaching pressures
mup = 0;
kinstruc_c3_00 = kinship_function(double(subs(subs(Umatc3,X,c3_Xsolve(:,:,1)))),Fmatc3);
related_high_c3_00 = relatedness_high_function(kinstruc_c3_00);
related_low_c3_00 = relatedness_low_function(kinstruc_c3_00);

mup = 0.05;
kinstruc_c3_05 = kinship_function(double(subs(subs(Umatc3,X,c3_Xsolve(:,:,26)))),Fmatc3);
related_high_c3_05 = relatedness_high_function(kinstruc_c3_05);
related_low_c3_05 = relatedness_low_function(kinstruc_c3_05);

mup = 0.10;
kinstruc_c3_10 = kinship_function(double(subs(subs(Umatc3,X,c3_Xsolve(:,:,51)))),Fmatc3);
related_high_c3_10 = relatedness_high_function(kinstruc_c3_10);
related_low_c3_10 = relatedness_low_function(kinstruc_c3_10);

mup = 0.20;
kinstruc_c3_20 = kinship_function(double(subs(subs(Umatc3,X,c3_Xsolve(:,:,101)))),Fmatc3);
related_high_c3_20 = relatedness_high_function(kinstruc_c3_20);
related_low_c3_20 = relatedness_low_function(kinstruc_c3_20);

% The following steps calculate the expected oldest age for all ages of
% focal under the different poaching pressures.

% make empty vectors for results
Py_c0_c00 = zeros(63,64);
Py_c0_c05 = zeros(63,64);
Py_c0_c10 = zeros(63,64);
Py_c0_c20 = zeros(63,64);
Py_c3_c00 = zeros(63,64);
Py_c3_c05 = zeros(63,64);
Py_c3_c10 = zeros(63,64);
Py_c3_c20 = zeros(63,64);

% calculate total age structure of family
z_c0_00 = sum(kinstruc_c0_00(:,1:64,2:end),3);
z_c0_05 = sum(kinstruc_c0_05(:,1:64,2:end),3);
z_c0_10 = sum(kinstruc_c0_10(:,1:64,2:end),3);
z_c0_20 = sum(kinstruc_c0_20(:,1:64,2:end),3);
z_c3_00 = sum(kinstruc_c3_00(:,1:64,2:end),3);
z_c3_05 = sum(kinstruc_c3_05(:,1:64,2:end),3);
z_c3_10 = sum(kinstruc_c3_10(:,1:64,2:end),3);
z_c3_20 = sum(kinstruc_c3_20(:,1:64,2:end),3);

% calculate probability that oldest age is 1 at birth of focal
Py_c0_00(1,1) = ( z_c0_00(1,1) / sum(z_c0_00(:,1)))^sum(z_c0_00(:,1));
Py_c0_05(1,1) = ( z_c0_05(1,1) / sum(z_c0_05(:,1)))^sum(z_c0_05(:,1));
Py_c0_10(1,1) = ( z_c0_10(1,1) / sum(z_c0_10(:,1)))^sum(z_c0_10(:,1));
Py_c0_20(1,1) = ( z_c0_20(1,1) / sum(z_c0_20(:,1)))^sum(z_c0_20(:,1));
Py_c3_00(1,1) = ( z_c3_00(1,1) / sum(z_c3_00(:,1)))^sum(z_c3_00(:,1));
Py_c3_05(1,1) = ( z_c3_05(1,1) / sum(z_c3_05(:,1)))^sum(z_c3_05(:,1));
Py_c3_10(1,1) = ( z_c3_10(1,1) / sum(z_c3_10(:,1)))^sum(z_c3_10(:,1));
Py_c3_20(1,1) = ( z_c3_20(1,1) / sum(z_c3_20(:,1)))^sum(z_c3_20(:,1));

% Calculate probability that i is oldest age at birth of focal
for i = 2:63
    Py_c0_00(i,1) = (sum(z_c0_00(1:i,1)) / sum(z_c0_00(:,1)))^sum(z_c0_00(:,1)) - (sum(z_c0_00(1:i-1,1)) / sum(z_c0_00(:,1)))^sum(z_c0_00(:,1));
    Py_c0_05(i,1) = (sum(z_c0_05(1:i,1)) / sum(z_c0_05(:,1)))^sum(z_c0_05(:,1)) - (sum(z_c0_05(1:i-1,1)) / sum(z_c0_05(:,1)))^sum(z_c0_05(:,1));
    Py_c0_10(i,1) = (sum(z_c0_10(1:i,1)) / sum(z_c0_10(:,1)))^sum(z_c0_10(:,1)) - (sum(z_c0_10(1:i-1,1)) / sum(z_c0_10(:,1)))^sum(z_c0_10(:,1));
    Py_c0_20(i,1) = (sum(z_c0_20(1:i,1)) / sum(z_c0_20(:,1)))^sum(z_c0_20(:,1)) - (sum(z_c0_20(1:i-1,1)) / sum(z_c0_20(:,1)))^sum(z_c0_20(:,1));
    Py_c3_00(i,1) = (sum(z_c3_00(1:i,1)) / sum(z_c3_00(:,1)))^sum(z_c3_00(:,1)) - (sum(z_c3_00(1:i-1,1)) / sum(z_c3_00(:,1)))^sum(z_c3_00(:,1));
    Py_c3_05(i,1) = (sum(z_c3_05(1:i,1)) / sum(z_c3_05(:,1)))^sum(z_c3_05(:,1)) - (sum(z_c3_05(1:i-1,1)) / sum(z_c3_05(:,1)))^sum(z_c3_05(:,1));
    Py_c3_10(i,1) = (sum(z_c3_10(1:i,1)) / sum(z_c3_10(:,1)))^sum(z_c3_10(:,1)) - (sum(z_c3_10(1:i-1,1)) / sum(z_c3_10(:,1)))^sum(z_c3_10(:,1));
    Py_c3_20(i,1) = (sum(z_c3_20(1:i,1)) / sum(z_c3_20(:,1)))^sum(z_c3_20(:,1)) - (sum(z_c3_20(1:i-1,1)) / sum(z_c3_20(:,1)))^sum(z_c3_20(:,1));
end

% calculate probability that age i is the oldest age at age x-1 of focal
for x = 1:62
    % this is the case that i=x and focal is the oldest individual
    Py_c0_00(x,x+1) = ( z_c0_00(x,x+1) / sum(z_c0_00(x:end,x+1)))^sum(z_c0_00(x:end,x+1));
    Py_c0_05(x,x+1) = ( z_c0_05(x,x+1) / sum(z_c0_05(x:end,x+1)))^sum(z_c0_05(x:end,x+1));
    Py_c0_10(x,x+1) = ( z_c0_10(x,x+1) / sum(z_c0_10(x:end,x+1)))^sum(z_c0_10(x:end,x+1));
    Py_c0_20(x,x+1) = ( z_c0_20(x,x+1) / sum(z_c0_20(x:end,x+1)))^sum(z_c0_20(x:end,x+1));
    Py_c3_00(x,x+1) = ( z_c3_00(x,x+1) / sum(z_c3_00(x:end,x+1)))^sum(z_c3_00(x:end,x+1));
    Py_c3_05(x,x+1) = ( z_c3_05(x,x+1) / sum(z_c3_05(x:end,x+1)))^sum(z_c3_05(x:end,x+1));
    Py_c3_10(x,x+1) = ( z_c3_10(x,x+1) / sum(z_c3_10(x:end,x+1)))^sum(z_c3_10(x:end,x+1));
    Py_c3_20(x,x+1) = ( z_c3_20(x,x+1) / sum(z_c3_20(x:end,x+1)))^sum(z_c3_20(x:end,x+1));
    
    % this loops through all cases i>x
    for i = (x+1):63
       Py_c0_00(i,x+1) = (sum(z_c0_00(x:i,x+1)) / sum(z_c0_00(x:end,x+1)))^sum(z_c0_00(x:end,x+1)) - (sum(z_c0_00(x:i-1,x+1)) / sum(z_c0_00(x:end,x+1)))^sum(z_c0_00(x:end,x+1));
       Py_c0_05(i,x+1) = (sum(z_c0_05(x:i,x+1)) / sum(z_c0_05(x:end,x+1)))^sum(z_c0_05(x:end,x+1)) - (sum(z_c0_05(x:i-1,x+1)) / sum(z_c0_05(x:end,x+1)))^sum(z_c0_05(x:end,x+1));
       Py_c0_10(i,x+1) = (sum(z_c0_10(x:i,x+1)) / sum(z_c0_10(x:end,x+1)))^sum(z_c0_10(x:end,x+1)) - (sum(z_c0_10(x:i-1,x+1)) / sum(z_c0_10(x:end,x+1)))^sum(z_c0_10(x:end,x+1));
       Py_c0_20(i,x+1) = (sum(z_c0_20(x:i,x+1)) / sum(z_c0_20(x:end,x+1)))^sum(z_c0_20(x:end,x+1)) - (sum(z_c0_20(x:i-1,x+1)) / sum(z_c0_20(x:end,x+1)))^sum(z_c0_20(x:end,x+1));
       Py_c3_00(i,x+1) = (sum(z_c3_00(x:i,x+1)) / sum(z_c3_00(x:end,x+1)))^sum(z_c3_00(x:end,x+1)) - (sum(z_c3_00(x:i-1,x+1)) / sum(z_c3_00(x:end,x+1)))^sum(z_c3_00(x:end,x+1));
       Py_c3_05(i,x+1) = (sum(z_c3_05(x:i,x+1)) / sum(z_c3_05(x:end,x+1)))^sum(z_c3_05(x:end,x+1)) - (sum(z_c3_05(x:i-1,x+1)) / sum(z_c3_05(x:end,x+1)))^sum(z_c3_05(x:end,x+1));
       Py_c3_10(i,x+1) = (sum(z_c3_10(x:i,x+1)) / sum(z_c3_10(x:end,x+1)))^sum(z_c3_10(x:end,x+1)) - (sum(z_c3_10(x:i-1,x+1)) / sum(z_c3_10(x:end,x+1)))^sum(z_c3_10(x:end,x+1));
       Py_c3_20(i,x+1) = (sum(z_c3_20(x:i,x+1)) / sum(z_c3_20(x:end,x+1)))^sum(z_c3_20(x:end,x+1)) - (sum(z_c3_20(x:i-1,x+1)) / sum(z_c3_20(x:end,x+1)))^sum(z_c3_20(x:end,x+1));

    end

end

% The probability that 63 is the oldest age is focal is 63 is always one.
Py_c0_00(63,64) = 1;
Py_c0_05(63,64) = 1;
Py_c0_10(63,64) = 1;
Py_c0_20(63,64) = 1;
Py_c3_00(63,64) = 1;
Py_c3_05(63,64) = 1;
Py_c3_10(63,64) = 1;
Py_c3_20(63,64) = 1;

% calculate expected max age from probabilities
maxage_c0_00 = ((1:63)*Py_c0_00)';
maxage_c0_05 = ((1:63)*Py_c0_05)';
maxage_c0_10 = ((1:63)*Py_c0_10)';
maxage_c0_20 = ((1:63)*Py_c0_20)';
maxage_c3_00 = ((1:63)*Py_c3_00)';
maxage_c3_05 = ((1:63)*Py_c3_05)';
maxage_c3_10 = ((1:63)*Py_c3_10)';
maxage_c3_20 = ((1:63)*Py_c3_20)';

% create figure
c3_maxage_fig = figure('name','c3_maxage_fig');

tiledlayout(1,3)

% mup = 0.05
nexttile
line([0 63],[0 63],'Color','black')
hold on
plot(0:(length(maxage_c0_00)-1), maxage_c0_00, c0_0_style )
plot(0:(length(maxage_c0_05)-1), maxage_c0_05, 'Color', c0_colour)
plot(0:(length(maxage_c3_05)-1), maxage_c3_05, 'Color', c3_colour)
hold off 
xlim([0 63])
legend("",'No poaching','No feedback','Effect of matriarch age on juvenile survival')
xlabel('Age of Focal (x)')
ylabel('Matriarch age')
title("Low poaching (\mu = 0.05)")

% mup = 0.1
nexttile
line([0 63],[0 63],'Color','black')
hold on
plot(0:(length(maxage_c0_00)-1), maxage_c0_00, c0_0_style )
plot(0:(length(maxage_c0_10)-1), maxage_c0_10, 'Color', c0_colour)
plot(0:(length(maxage_c3_10)-1), maxage_c3_10, 'Color', c3_colour)
hold off 
xlim([0 63])
legend("",'No poaching','No feedback','Effect of matriarch age on juvenile survival')
xlabel('Age of Focal (x)')
ylabel('Matriarch age')
title("Medium poaching (\mu = 0.10)")

% mup = 0.2
nexttile
line([0 63],[0 63],'Color','black')
hold on
plot(0:(length(maxage_c0_00)-1), maxage_c0_00, c0_0_style )
plot(0:(length(maxage_c0_20)-1), maxage_c0_20, 'Color', c0_colour)
plot(0:(length(maxage_c3_20)-1), maxage_c3_20, 'Color', c3_colour)
hold off 
xlim([0 63])
legend("",'No poaching','No feedback','Effect of matriarch age on juvenile survival')
xlabel('Age of Focal (x)')
ylabel('Matriarch age')
title("High poaching (\mu = 0.20)")

% set figure size
c3_maxage_fig.Units = 'centimeters';
c3_maxage_fig.Position = [0 0 54 18];

% save figure
exportgraphics(c3_maxage_fig,'fig7.pdf','Resolution',600)


%% PLOT MATRIARCH AGE ON SURVIVAL FULL KINSTRUCTURE
% Here the full kin sturcture with the effect of matriarch age on survival
% Figure S4

% rotate kinstructure
total_kinstruc_c3_05 = shiftdim(sum(kinstruc_c3_05),1);
total_kinstruc_c3_10 = shiftdim(sum(kinstruc_c3_10),1);
total_kinstruc_c3_20 = shiftdim(sum(kinstruc_c3_20),1);

% create figure
c3_totkin_fig = figure('name','c3_totkin_fig');

tiledlayout(3,3)

% mup = 0.05 old generation
nexttile
area([total_kinstruc_c3_05(:,[5 6 7]) sum(total_kinstruc_c3_05(:,[12 13]),2) ] ) 
xlim([0 63])
ylim([0 2])
legend('Mother','Grandmother','Great-grandmother','Aunts')
xlabel('Age of Focal (x)')
ylabel('Number')
title("Low poaching (\mu = 0.05)")

% mup = 0.1 old generation
nexttile
area([total_kinstruc_c3_10(:,[5 6 7]) sum(total_kinstruc_c3_10(:,[12 13]),2) ] ) 
xlim([0 63])
ylim([0 2])
legend('Mother','Grandmother','Great-grandmother','Aunts')
xlabel('Age of Focal (x)')
ylabel('Number')
title("Medium poaching (\mu = 0.10)")

% mup = 0.2 old generation
nexttile
area([total_kinstruc_c3_20(:,[5 6 7]) sum(total_kinstruc_c3_20(:,[12 13]),2) ] ) 
xlim([0 63])
ylim([0 2])
legend('Mother','Grandmother','Great-grandmother','Aunts')
xlabel('Age of Focal (x)')
ylabel('Number')
title("High poaching (\mu = 0.20)")

% mup = 0.05 middle generation
nexttile
area([sum(total_kinstruc_c3_05(:,[8 9]),2) sum(total_kinstruc_c3_05(:,[14 15]),2) ] ) 
xlim([0 63])
ylim([0 1.5])
legend('Sisters','Causins')
xlabel('Age of Focal (x)')
ylabel('Number')
title("Low poaching (\mu = 0.05)")

% mup = 0.1 middle generation
nexttile
area([sum(total_kinstruc_c3_10(:,[8 9]),2) sum(total_kinstruc_c3_10(:,[14 15]),2) ] ) 
xlim([0 63])
ylim([0 1.5])
legend('Sisters','Causins')
xlabel('Age of Focal (x)')
ylabel('Number')
title("Medium poaching (\mu = 0.10)")

% mup = 0.2 middle generation
nexttile
area([sum(total_kinstruc_c3_20(:,[8 9]),2) sum(total_kinstruc_c3_20(:,[14 15]),2) ] ) 
xlim([0 63])
ylim([0 1.5])
legend('Sisters','Causins')
xlabel('Age of Focal (x)')
ylabel('Number')
title("High poaching (\mu = 0.20)")

% mup = 0.05 young generation
nexttile
area([total_kinstruc_c3_05(:,[2 3 4]) sum(total_kinstruc_c3_05(:,[10 11]),2) ] ) 
xlim([0 63])
ylim([0 3])
legend('Daughters','Granddaughters', 'Great-granddaughters',"Nieces",'Location','northwest')
xlabel('Age of Focal (x)')
ylabel('Number')
title("Low poaching (\mu = 0.05)")

% mup = 0.1 young generation
nexttile
area([total_kinstruc_c3_10(:,[2 3 4]) sum(total_kinstruc_c3_10(:,[10 11]),2) ] ) 
xlim([0 63])
ylim([0 3])
legend('Daughters','Granddaughters', 'Great-granddaughters',"Nieces",'Location','northwest')
xlabel('Age of Focal (x)')
ylabel('Number')
title("Medium poaching (\mu = 0.10)")

% mup = 0.2 young generation
nexttile
area([total_kinstruc_c3_20(:,[2 3 4]) sum(total_kinstruc_c3_20(:,[10 11]),2) ] ) 
xlim([0 63])
ylim([0 3])
legend('Daughters','Granddaughters', 'Great-granddaughters',"Nieces",'Location','northwest')
xlabel('Age of Focal (x)')
ylabel('Number')
title("High poaching (\mu = 0.20)")

% set figure size
c3_totkin_fig.Units = 'centimeters';
c3_totkin_fig.Position = [0 0 54 70];

% size figure
exportgraphics(c3_totkin_fig,'c3_totkin_fig.pdf','Resolution',600)


%% ALL FEEDBACK COMBINED (c4)

% reset values
syms mup;
Fmatc4 = sym(Fmat0);
Umatc4 = Umat0;
X = sym('x',[1, length(c4Xbase)]);


% calculate baseline values without poaching
c4Xbase = c4_calcX(kinstruc_base);

% include feedback in matrices
Umatc4(:,1:18) = (1-(c4Xbase(1:18) - X(1:18)).*alphax(1:18)').*Umat0(:,1:18);
Fmatc4(:,9:20) = (1 + ( X(19:30) - c4Xbase(19:30) ) .* kappa) .* Fmat0(:,9:20);
Umatc4(:,1:8) = Umatc4(:,1:8) .* (1+exp(0.047.*c4Xbase(31:38))) ./ (1+exp(0.047.*X(31:38))) .* exp(0.047*(X(31:38)-c4Xbase(31:38)));

% make empty arrays for results
c4_lambdas = zeros(1,101);
c4_eigvec = zeros(63,101);
c4_Xsolve = zeros(1,length(c4Xbase),101);
c4_poprel_low = zeros(1,101);
c4_poprel_high = zeros(1,101);

% set values without poaching
c4_lambdas(1) = c0_lambdas(1);
c4_Xsolve(:,:,1) = c4Xbase;
c4_eigvec(:,1) = c0_eigvec(:,1);
c4_poprel_low(1) = c0_poprel_low(1);
c4_poprel_high(1) = c0_poprel_high(1);

% iterate through all poaching pressures
for i = 2:length(mupvalues)
    
    % set poaching
    mup = mupvalues(i);
    
    % solve kinstructure
    c4_Xsolve(:,:,i) = fsolve(@(X)solveXkin(X,Fmatc4,subs(Umatc4),@c4_calcX),c4_Xsolve(:,:,i-1),optimoptions('fsolve','Display','iter','UseParallel',false));

    % create population projection matrix
    Amatc4 = double(subs(subs(Umatc4,X,c4_Xsolve(:,:,i))) + subs(subs(Fmatc4,X,c4_Xsolve(:,:,i))));
    
    % calculate population growth rate and structure
    [c4_eigvec(:,i), c4_lambdas(i)]  = eigs(Amatc4,1);
    c4_eigvec(:,i) = c4_eigvec(:,i)/sum(c4_eigvec(:,i));

    % calculate full kinstructure
    kinstruc_temp = kinship_function( double(subs(subs(Umatc4,X,c4_Xsolve(:,:,i)))),double(subs(subs(Fmatc4,X,c4_Xsolve(:,:,i)))));
    
    % calculate relatedness
    related_high_temp = relatedness_high_function(kinstruc_temp);
    related_low_temp = relatedness_low_function(kinstruc_temp);
    
    c4_poprel_low(i) = related_low_temp(1:63)*c4_eigvec(:,i);
    c4_poprel_high(i) = related_high_temp(1:63)*c4_eigvec(:,i);

    % report progress
    fprintf('\n\n\nSucceeded, mup=%f, lambda = %f, progress %f\n', mup, c4_lambdas(i), i/101);
end


%% PLOT POPULATION GROWTH RATES
% this creates the figure with the growth rate under the range of poaching
% presures with all types of feedback.
% Figure 2

% create figure
popgrowth_fig = figure('name','popgrowth_fig');

yline(1)
xlabel('Poaching pressure (\mu)')
ylabel('population growthrate (\lambda)')
hold on
plot(mupvalues, c0_lambdas, 'color',c0_colour)
plot(mupvalues, c1_lambdas, 'color',c1_colour)
plot(mupvalues, c2_lambdas, 'color',c2_colour)
plot(mupvalues, c3_lambdas, 'color',c3_colour)
plot(mupvalues, c4_lambdas, 'color',c4_colour)
hold off
legend("",'No feedback','Effect of mothers on survival','Effect of sisters on fecundity','Effect of matriarch age on juvenile survival','All effects combined','location','southwest')
ylim([0.80, 1.05])

% set figure size
popgrowth_fig.Units = 'centimeters';
popgrowth_fig.Position = [0 0 18 18];

% save figure
exportgraphics(popgrowth_fig,'fig2.pdf','Resolution',600)


%% PLOT RELATEDNESS
% This section plots the expected relatedness between two individuals in
% the population under a range of poaching conditions for all family
% interactions.
% Figure 4

% create figure
poprel_fig = figure('name','poprel_fig');
xlabel('Poaching pressure (\mu)')
ylabel('Expected relatedness (\xi)')
hold on
plot(mupvalues, c0_poprel_low,'--', 'color',c0_colour)
plot(mupvalues, c1_poprel_low,'--', 'color',c1_colour)
plot(mupvalues, c2_poprel_low,'--', 'color',c2_colour)
plot(mupvalues, c3_poprel_low,'--', 'color',c3_colour)
plot(mupvalues, c4_poprel_low,'--', 'color',c4_colour)
plot(mupvalues, c0_poprel_high,'-', 'color',c0_colour)
plot(mupvalues, c1_poprel_high,'-', 'color',c1_colour)
plot(mupvalues, c2_poprel_high,'-', 'color',c2_colour)
plot(mupvalues, c3_poprel_high,'-', 'color',c3_colour)
plot(mupvalues, c4_poprel_high,'-', 'color',c4_colour)
hold off
legend('','','','','','No feedback','Effect of mothers on survival','Effect of sisters on fecundity','Effect of matriarch age on juvenile survival','All effects combined','location','southeast')
ylim([0 0.5])

% set figure size
popgrowth_fig.Units = 'centimeters';
popgrowth_fig.Position = [0 0 18 18];

% save figure
exportgraphics(poprel_fig,'fig4.pdf','Resolution',600)


%% CLEANUP AND SAVE WORKSPACE

% close all figures
close all

% save workspace with calculated values
save('elephantworkspace')
```
