## Supplementary material for "Family matters: Linking population growth, kin interactions, and African elephant social groups": Matlab code to run the model: README.rtf

GENERAL INFORMATION
This README.txt file was updated on 11 March 2024

A. Paper associated with this archive
Citation:
Family matters: Linking population growth, kin interactions, and African elephant social groups
Brief abstract: In many species, individuals are embedded in a network of kin with whom they interact. The interactions among kin may affect the survival and fertility rates, and thus the life history of individuals. These interactions indirectly influence both the network of kin and the dynamics of the population. In this way, non-linear feedback emerges between the kin network and individual life history rates. We describe a model that calculates the kin network of an individual while incorporating the feedback between the network and the life history of the individual. We demonstrate the use of this model for African elephant populations. We incorporate effects of the mother's presence and matriarch age on the survival of juveniles, and of the presence of a sister on the fecundity of young females.  We find that interactions between family members amplify the negative effects of poaching on the family structure and growth rate of African elephant populations. Our analysis provides a framework that can be applied to a broad range of of social species.

B. Originators
Names, institutions of all authors

C. Contact information
Name
Address
email

F. Funding Sources
[list all sources of funding that supported the collection of the data here]
OMIT this General Information (above) for double-blind review but include it on final acceptance

ACCESS INFORMATION
1. Licenses/restrictions placed on the data or code
CC BY (Attribution)

3. Recommended citation for this data/code archive
Cite with article citation.

DATA & CODE FILE OVERVIEW
This data repository consist of 9 code scripts, and this README document, with the following data and code filenames:

Code scripts and workflow
[file names and brief descriptions. Also describe the workflow if there are several scripts that need to be run in order]
    	1. Main_elephant_kin_interactions.m; This is the main script that is used for analysis of computations and graphs. All other files contain MATLAB functions called from this file.
	2. kinship_function.m; This file contains the kinship_function(Umat, Fmat) function, which takes the survival matrix (Umat) and fecundity matrix (Fmat) and returns a array with the kinship structure calculated according to the equations in the main text of the paper.
	3. relatedness_high_function.m; This file contains the relatedness_high_function(kinstruc) function, which takes a kinship structure as produced by kinship_function.m and returns the higher estimate of the relatedness of Focal to the kin network calculated according to the equations in the main text of the paper.
	4. relatedness_low_function.m; This file contains the relatedness_low_function(kinstruc) function, which takes a kinship structure as produced by kinship_function.m and returns the lower estimate of the relatedness of Focal to the kin network calculated according to the equations in the main text of the paper.
	5. c1_calcX.m; This file contains the c1_calcX(kinstruc) function, which takes a kinship structure as produced by kinship_function.m and returns a vector with the age specific probability an individual has a living mother, to implement the feedback between mother presence and juvenile survival as discussed in the main text of the paper. 
	6. c2_calcX.m; This file contains the c2_calcX(kinstruc) function, which takes a kinship structure as produced by kinship_function.m and returns a vector with the age specific probability of having at least one sister, to implement the feedback between the presence of a sister and fecundity as discussed in the main text of the paper. 
	7. c3_calcX.m; This file contains the c3_calcX(kinstruc) function, which takes a kinship structure as produced by kinship_function.m and returns a vector with the expected oldest age in the kinship network, to implement the feedback between matriarch age and juvenile survival as discussed in the main text of the paper. 
	8. c4_calcX.m; This file contains the c4_calcX(kinstruc) function, is a wrapper combining the c1_calcX.m, c2_calcX.m and c3_calcX.m to implement the feedback effects from these functions simultaneously.
	9. solveXkin.m; This file contains the solveXkin(X,Fmat,Umat,@calcX), which is a wrapper of the c1_calcX.m, c2_calcX.m, c3_calcX.m and c4_calcX.m functions to feed these functions in the solve function and in this way solve the kinship network assuming these interactions. The function takes a guess of the values produced by the c1_calcX.m, c2_calcX.m, c3_calcX.m and c4_calcX.m functions (X), the fecundity matrix (Fmat), the survival matrix (Umat) and a handle of the c1_calcX.m, c2_calcX.m, c3_calcX.m and c4_calcX.m functions. The function returns the difference between the provided guess values (X) and the values calculated based on the provided matrices and function handle. 


SOFTWARE VERSIONS
MATLAB version 9.14.0.2254940 (R2023a) Update 2
Matworks Optimisation Toolbox version 9.5
