## Supplementary material for "Family matters: Linking population growth, kin interactions, and African elephant social groups": Matlab code to run the model: relatedness_high_function.m

```
function relatedness = relatedness_high_function(allkin)
% This function calculates the expected relatedness of focal to the
% provided kinship network assuming that all sisters share the same father.
%
% allkin = array with the age specific kin network as generated by the
% kinship function in kinship_function.m
%
% the output is a array with the age specific relatedness

% the relatedness to all types of kin in the kin network
reatedness_value = [0 1/2 1/4 1/8 1/2 1/4 1/8 1/2 1/2 1/4 1/4 1/4 1/4 1/8 1/8]';

% collapse the age structure of kin
kinsum = permute(sum(allkin),[3,2,1]);

% calculate expected relatedness.
relatedness = reatedness_value(2:end, :)' * kinsum(2:end, :) ./ sum( kinsum(2:end,:) ); 

end
```
