## Supplementary material for "Family matters: Linking population growth, kin interactions, and African elephant social groups": Matlab code to run the model: solveXkin.m

```
function diff=solveXkin(Xguess, Fmatc, Umatc, Xcalc)
% This is a general wrapper function that calculates the difference between
% the guess of a set of unknown variables and the value of these values
% predicted by the kinship network functions using the Xcalc function. This
% difference is minimized by the fsolve function from matlab to solve the
% kinship network. In this way any feedback with any property of the
% kinship network can be entered in the fsolve function.
%
% Xguess = the current guess of the parameters
% Fmatc = fecundity matrix in which the entries might depend on the values in
% Xguess
% Umatc = survival matrix in which the entries might depend on the values in
% Xguess
% Xcalc = hadle (@) to a function that calculates the property of the
% kinship network presented by X.
%
% The output is the differnece between the Xguess and the values of X
% calculated from the kin network.

% make a dummy vector X
X = sym('x',[1 length(Xguess)]);

% substitute the Xguess values in the matrices
Fmatc = double(subs(Fmatc, X, Xguess));
Umatc = double(subs(Umatc, X, Xguess));

% calculate the kin network using the guessed values
kinstruc = kinship_function(Umatc,Fmatc);

% calculate the values represented by X from the kin network
Xvalue = Xcalc(kinstruc);

% calculate the difference between the calculated and guessed variables
diff = Xguess - Xvalue;

end
```
